# Pathologic α-Synuclein-NOD2 Interaction and RIPK2 Activation Drives Microglia-Induced Neuroinflammation in Parkinson’s Disease

**DOI:** 10.1101/2024.02.19.580982

**Authors:** Bo Am Seo, Seung-Hwan Kwon, Donghoon Kim, Han-Byeol Kim, Shi-Xun Ma, Kundlik Gadhave, Noelle Burgess, Xiaobo Mao, Liana S. Rosenthal, Javier Redding-Ochoa, Juan C Troncoso, Seulki Lee, Valina L. Dawson, Ted M. Dawson, Han Seok Ko

## Abstract

Pathological aggregation of α-Synuclein (α-Syn) and neuroinflammation are closely linked to Parkinson’s disease (PD). However, the specific regulators of the neuroinflammation caused by pathological α-syn remain obscure. In this study, we show that NOD2/RIPK2 signaling is a crucial regulator of neuroinflammation in PD. Pathological α-syn binds to NOD2, causing self-oligomerization and complex formation with RIPK2, leading to RIPK2 ubiquitination and activation of MAPK and NF-kB. Notably, this NOD2/RIPK2 signaling is particularly active in microglia of human PD brains and the α-Syn preformed fibril (α-Syn PFF) mouse model. Depleting NOD2 or RIPK2 reduces neuroinflammation and protects against dopamine neuron degeneration in a pathologic α-Syn mouse model by blocking the formation of neurotoxic reactive astrocytes caused by microglia activation. The discovery of NOD2/RIPK2 signaling as a key regulator of neuroinflammation in PD provides a new understanding of α-Syn-driven neuroinflammation and neurodegeneration in PD and a potential new therapeutic strategy.

**Graphical Abstract:** 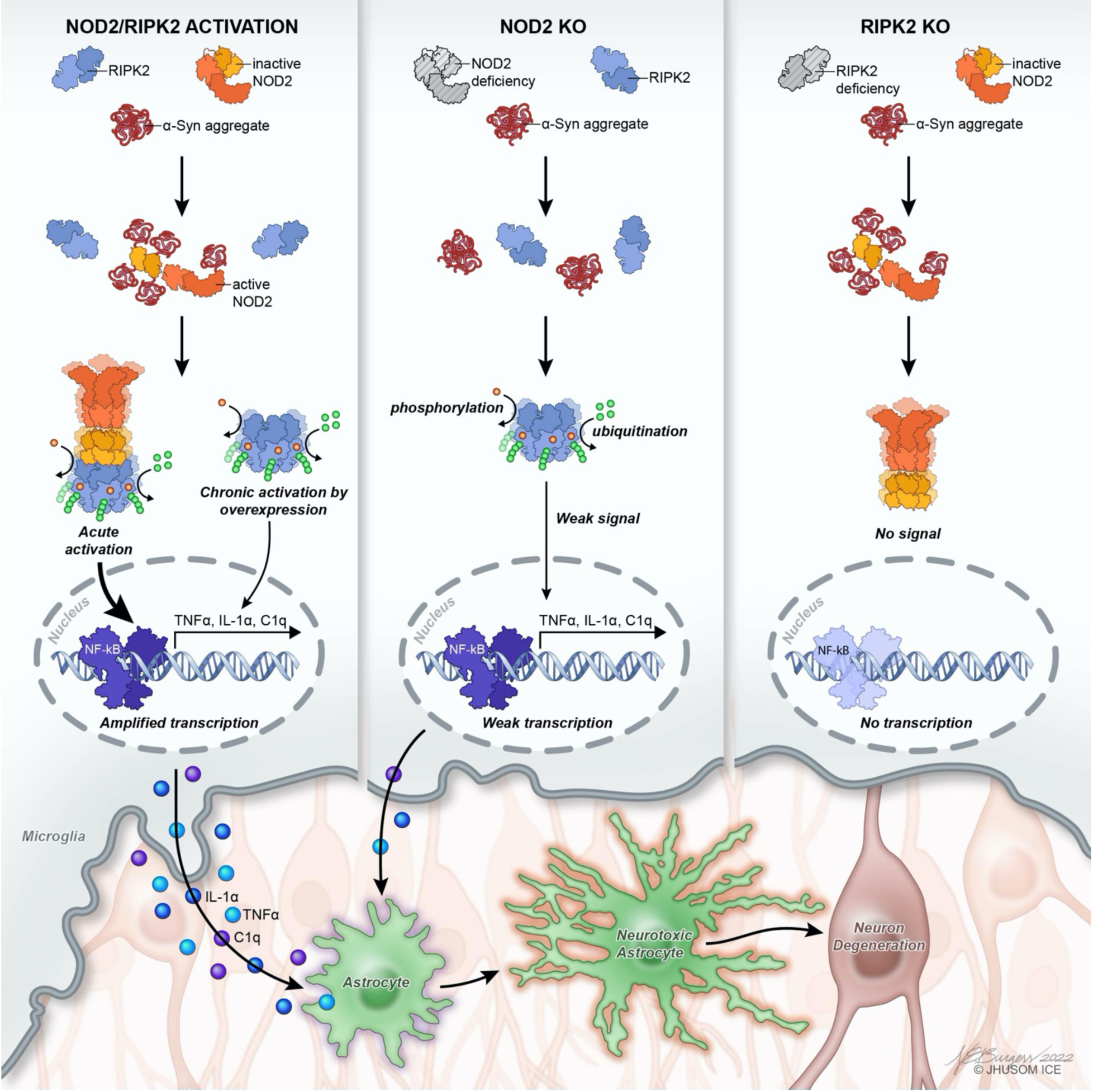

**In brief:** Pathological α-Synuclein (α-Syn) binds to the microglial NOD2 protein, which in turn triggers NOD2/RIPK2 complex and RIPK2 phosphorylation/ubiquitination. This process activates the NF-kB/MAPK pathways, ultimately leading to neurotoxic reactive astrocyte-induced dopaminergic neurodegeneration. Depletion of RIPK2 (RIPK2 KO) or NOD2 (NOD2) protects dopamine neurons in a mouse model of Parkinson’s disease (PD). These findings provide insights into α-Syn-driven neuroinflammation and offer potential therapeutic strategies for PD.

**Highlights:**

NOD2/RIPK2 signaling is identified as a crucial regulator of neuroinflammation in PD.
NOD2/RIPK2 signaling is highly active in microglia in human PD brains and α-Syn PFF mouse models.
Pathological α-Syn binds to NOD2, triggering self-oligomerization and RIPK2 complex formation, leading to MAPK and NF-kB activation
Genetic depletion of NOD2 or RIPK2 reduces neuroinflammation and protects dopamine neurons by blocking the formation of neurotoxic reactive astrocytes.

## Introduction

Parkinson’s disease (PD), a neurodegenerative disorder, is characterized by the progressive loss of dopamine (DA) neurons in the substantia nigra pars compacta (SNc). A defining feature of PD is the formation of Lewy bodies (LBs), intracellular accumulations of α-syn. Pathological α-syn has a strong correlation with the onset and progression of PD, as it drives the disease’s pathogenesis and neurodegeneration through both cell-autonomous and non-cell-autonomous mechanisms. Accumulation of pathological α-syn leads to neurodegeneration in PD, through multiple mechanisms including disruptions to synaptic vesicle trafficking, mitochondrial, endoplasmic reticulum, and Golgi function, suppression of the endocytic, autophagic, or lysosomal pathways, misregulation of organelle dynamics, and interference with inter-organelle contacts and protein degradation processes (Michel et al., 2016; Wong and Krainc, 2017). Ultimately, these mechanisms initiated by pathologic α-syn act in a cell-autonomous manner within neurons to kill neurons through parthanatos (Kam et al., 2018; Park et al., 2022). However, recent research has also suggested that pathological α-syn contributes to neurodegeneration through non-cell-autonomous mechanisms involving glial cells in PD (Chou et al., 2021a; Lim et al., 2018; Yun et al., 2018).

α-Syn is a presynaptic protein that is abundant in the cytoplasm of neurons. There is increasing evidence to suggest that in response to cellular stress and injury, α-syn can be released by neurons into the extracellular space through exocytosis (Chou et al., 2021b; Lim et al., 2018; Rodriguez et al., 2018). This secreted α-syn can then be transmitted from one cell to another in a prion-like fashion (Ma et al., 2019). There are various mechanisms by which secreted α-syn can be transmitted to neighboring neurons, including unconventional endocytosis, tunneling nanotubes (TNTs), and receptor-mediated endocytosis (Abounit et al., 2016; Choi et al., 2021; Guo and Lee, 2014; Lagalwar, 2022). One of the receptors involved in this process is the lymphocyte activation gene 3 (LAG3). This intercellular transmission of α-syn leads to neuronal toxicity and the spread of pathological α-syn, resulting in the formation of Lewy-like inclusions (Gu et al., 2021; Mao et al., 2016; Zhang et al., 2023a; Zhang et al., 2023b; Zhang et al., 2021). This can also impact neighboring glial cells, leading to neuroinflammation in microglia and astrocytes. Neuroinflammation is increasingly recognized as a crucial factor in the pathogenesis of PD. Recent studies using human post-mortem tissue and animal models have provided strong evidence linking neuroinflammation to the onset and progression of PD (Hirsch and Standaert, 2021; Ma and Lim, 2021; Pajares et al., 2020; Smajic et al., 2022; Tansey et al., 2022; Wang et al., 2015; Yun et al., 2018). Neuroinflammation in PD is characterized by an increased presence of microglia and elevated levels of pro-inflammatory cytokines and chemokines (Ma and Lim, 2021; Smajic et al., 2022). Microglia are the key immune cells in the central nervous system (CNS), playing an important role in maintaining and supporting the neural network (Borst et al., 2021; Perry and Teeling, 2013). However, when microglia are chronically activated due to systemic inflammation or neurodegeneration, they can create a pro-inflammatory environment in the brain and contribute to the progression of neurodegeneration in PD (Garcia et al., 2022; George et al., 2019; Kam et al., 2020; Subramaniam and Federoff, 2017). We and others have recently established that pathological α-syn triggers the pro-inflammatory activation of microglia. Our findings have demonstrated that activation of microglia caused by pathological α-syn results in the secretion of IL-1α, TNFα, and C1q, which are inducers of neurotoxic reactive astrocytes that contribute to neuronal toxicity (Guttenplan et al., 2021; Liddelow et al., 2017; Yun et al., 2018). Evidence supporting this link between microglial activation, reactive astrocyte formation, and neurobehavioral deficits in PD was observed in mouse models of pathologic α-syn models of neurodegeneration (Yun et al., 2018).

Microglia and astrocytes, as well as neuroinflammation, have garnered considerable attention as critical factors in the pathophysiology of Parkinson’s disease (PD). However, the specific cellular and molecular mechanisms through which they participate in neuroinflammatory processes following exposure to pathological α-synuclein are poorly understood. To uncover the cellular mechanisms underlying microglial activation induced by pathologic α-syn, we performed RNA sequencing analysis using primary murine microglia cultures that were activated by pathologic α-syn preformed fibrils (PFF). Our screening efforts revealed the involvement of a previously unknown mediator of microglia activation in PD, the NOD2 (Nucleotide Binding Oligomerization Domain Containing 2)/RIPK2 (Receptor-Interacting Protein Kinase 2) complex. Genetic inhibition of NOD2/RIPK2 signaling prevented microglial activation and neurotoxic reactive astrocyte formation and neurodegeneration in models of PD. The involvement of NOD2/RIPK2 signaling in models of PD, coupled with the findings that NOD2/RIPK2 signaling is overactive in human PD brain samples provides a potential important new disease modifying therapeutic opportunity in PD.

## Results

### *NOD2/RIPK2* transcripts are upregulated in α-Syn PFF-treated microglia

To explore the cellular components responsible for microglia activation in response to pathologic α-syn, RNAseq was performed using primary cultured microglia treated with endotoxin free α-Syn PFF or PBS (Figure S1A). Comprehensive analysis of these data revealed 462 differentially expressed genes (DEGs) in α-Syn PFF-activated microglia compared to non-activated microglia (Figure S1B and S1C and Table S1). Notably, within the top-ranked DEGs, the receptor*-interacting serine/threonine-protein kinase 2 (RIPK2) and nucleotide-binding oligomerization domain containing protein 2 (NOD2),* a well-known upstream activator of RIPK2 (Philpott et al., 2014), exhibited a significant increase in α-Syn PFF-treated microglia (Figure S1B). To gain further insights into the biological processes, cellular components, and molecular functions associated with the DEGs induced by α-Syn PFF treatment, we performed a gene ontology (GO) and Kyoto Encyclopedia of Genes and Genomes (KEGG) pathway analysis (Ashburner et al., 2000). The results of the GO analysis revealed that the DEGs were highly enriched in biological processes related to the immune system process and response (Figure S1D). Moreover, KEGG analysis demonstrated the activation of the NOD-like receptor signaling pathway in microglia treated with α-Syn PFF, indicating the potential involvement of NOD2/RIPK2 signaling (Figure S1E). Subsequently, validation of the upregulated NOD2/RIPK2 transcripts and proteins induced by α-Syn PFF treatment in primary cultured microglia was performed by qPCR and western blot analysis (Figure S1A). A significant increase in the expression levels of NOD2 and RIPK2 mRNA (Figure S1F) and protein (Figure S1G and S1H) in response to α-Syn PFF treatment was observed (Figure S1F-S1H).

### Pathologic α-synuclein binds to NOD2

Based on the critical role of NOD2 in activating RIPK2 after a pathogenic infection (Philpott et al., 2014), we wondered whether NOD2 interacts with pathologic α-syn. An *in-situ* proximity ligation assay (PLA), a technology capable of detecting endogenous protein-protein interactions at the single molecule resolution inside the cell was utilized (Clausson et al., 2011; Seo et al., 2021; Soderberg et al., 2006) (Figure 1A). α-Syn PFF-treated WT primary microglia revealed numerous strong fluorescence signals using specific antibodies for pathologic α-syn and NOD2 protein indicating an interaction between pathologic α-syn and NOD2. No fluorescence signal was detected in α-syn PFF-treated NOD2-knockout microglia, providing evidence for the specificity of this interaction in microglia (Figure 1B and 1C).

**Figure 1.**
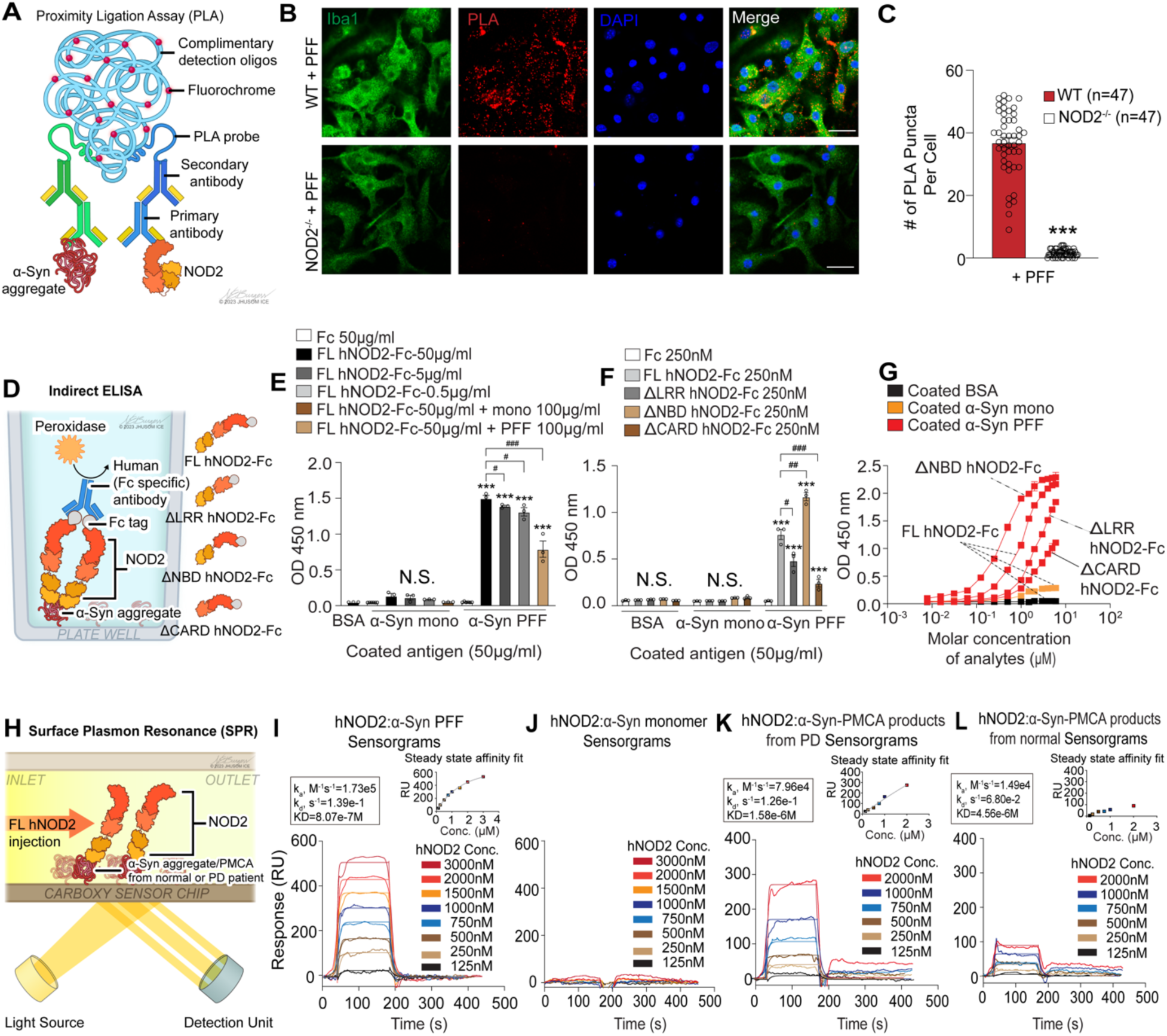
Pathologic α-Syn binds to NOD2 and activates RIPK2. (A) Schematic diagram illustrating PLA to visualize pathologic α-Syn aggregate interaction with endogenous NOD2 using α-Syn PFF-treated WT or NOD2^-/-^ microglia. (B) Confocal images showing PLA-positive signals (red dots) in α-Syn PFF-treated WT microglia, but not in NOD2^-/-^ microglia. (C) Bar graph depicting the number of red dots per cell in (B) (n=47 cells, biologically independent experiments). Data presented as mean ± SEM (***P < 0.001). (D) Schematic diagram of indirect ELISA confirming binding of pathologic α-Syn aggregates with hNOD2 protein using purified human IgG1 Fc-tagged human NOD2 full-length (FL hNOD2-Fc) and mutants lacking LRR, NBD, or CARD domain (ΔLRR hNOD2-Fc, ΔNBD hNOD2-Fc, and ΔCARD hNOD2-Fc). (E) Quantification of FL hNOD2-Fc binding to α-Syn PFF or α-Syn monomer using dose-dependent signal and competition ELISA (n=3, biologically independent). Data presented as mean ± SEM. Statistical significance: ***P < 0.001 (compared to Fc 50 μg/ml); ^#^P < 0.05, ^###^P < 0.001 (compared to FL hNOD2-Fc 50 μg/ml). (F) Quantification of FL hNOD2-Fc, ΔLRR hNOD2-Fc, ΔNBD hNOD2-Fc, or ΔCARD hNOD2-Fc binding to α-Syn PFF or α-Syn monomer (n=3, biologically independent). Data presented as mean ± SEM. Statistical significance: ***P < 0.001 (compared to Fc 250 nM); ^#^P < 0.05, ^##^P < 0.01, ^###^P < 0.001 (compared to FL hNOD2-Fc 250 nM). (G) Fitting curve for the binding affinity of FL hNOD2-Fc, ΔLRR hNOD2-Fc, ΔNBD hNOD2-Fc, or ΔCARD hNOD2-Fc to α-Syn PFF (n=3, biologically independent). The estimated affinity KD values of mutant forms with α-Syn PFF were 1.20 μM, 2.81 μM, 0.42 μM, and 16.42 μM. Binding affinity KD values determined by molar concentration at half-max OD value. Data presented as mean ± SEM. (H) Illustration of SPR experiments for real-time interaction measurement between α-Syn aggregate and hNOD2 protein. (I) The sensorgram and steady-state affinity fit showed FL hNOD2 binding to immobilized α-Syn PFF, along with the overlaid kinetics. Kinetic data was fitted to a 1:1 binding model, resulting in a KD of 8.07e-7M. Steady-state affinity calculation, aligned with kinetic data, yielded an affinity of 1.19e-6M (n=3, biologically independent SPR experiments). (J) The sensorgram showed FL hNOD2 not binding to immobilized α-Syn monomer. (n=3, biologically independent SPR experiments). (K) The sensorgram and steady-state affinity fit demonstrated FL hNOD2 binding to immobilized human PD patient-derived PMCA products. KD was calculated at 1.58e-6M (n=3, biologically independent SPR experiments). (L) The sensorgram and steady-state affinity fit revealed FL hNOD2 binding to immobilized healthy control-derived PMCA products. KD was calculated at 4.56e-6M (n=3, biologically independent SPR experiments).

To investigate the interaction of α-Syn PFF with human NOD2 protein (hNOD2), we performed an *in vitro* pull-down assay using previously well characterized biotin-conjugated α- Syn PFF (α-Syn-biotin PFF) and monomer (α-Syn-biotin monomer) (Mao et al., 2016). For this assay, HEK293T cells were transfected with plasmids expressing EGFP-fused hNOD2 and as controls other members of the intracellular NOD-like receptor protein (NLRs) family (Franchi et al., 2009), hNOD1 (a closely related protein with hNOD2) and hNLRP3 were used, and compared to hRIPK2 (Figure S2A-C). These transfected cells were treated with either α-Syn-biotin PFF or α-syn-biotin monomer, followed by lysis and pull-down using neutravidin (Figure S2A). α-Syn-biotin PFF binds only to hNOD2, while no such interaction was observed with α-syn-biotin monomer and α-syn-biotin PFF does not bind hNOD1, hNLRP3 and hRIPK2 (Figure S2B and S2C). NOD2 contains two caspase activation and recruitment domains (CARDs), a nucleotide-binding domain (NBD), and leucine-rich repeats (LRRs), which enable it to sense the structures of bacterial peptidoglycans, such as muramyl dipeptide (MDP) (Philpott et al., 2014). To elucidate the domain of hNOD2 that is responsible for binding with pathologic α-Syn, hNOD2 deletion mutants with His- and Myc-tag sequences, including hNOD2 mutant lacking the two CARD domains (ΔCARD), hNOD2 mutant lacking the NBD domain (ΔNBD), and hNOD2 mutant lacking the LRR domain (ΔLRR) were used (Sabbah et al., 2009). These mutants, along with full-length (FL) hNOD2, were expressed in HEK293T cells, followed by treatment with either α-Syn-biotin PFF or α-Syn-biotin monomer (Figure S2D). After the cells were lysed reciprocal pull-down assays were conducted using nickel-NTA agarose or neutravidin (Figure S2D). These pull-down assays revealed that α-Syn-biotin PFF, but not α-Syn-biotin monomer, specifically binds to the CARD domains of hNOD2 (Figure S2E and S2F).

To further investigate the interaction between hNOD2 and α-Syn PFF, an indirect enzyme-linked immunosorbent assay (ELISA) was utilized (Figure 1D). FL hNOD2 and its domain deletion mutants were purified using a human Fc-tag with HRV 3C (human rhinovirus 3C) cleavage sequence (Figure S2G) followed by an indirect ELISA with FL hNOD2 and mutants (Figure S2H-S2K) to validate the specific and direct binding between hNOD2 and α-Syn PFF. The results from the indirect ELISA confirmed the binding between α-Syn PFF and FL hNOD2 (Figure 1E). Adding α-Syn PFF (100 μg/ml) to the ELISA assay reduced the signal between α-Syn PFF and FL hNOD2 (Figure 1E). Minimal to no signal was detected between BSA or α-Syn monomer and FL hNOD2 (Figure 1E). The interaction signal significantly decreased when the CARD domains of hNOD2 were deleted while the deletion of the LRR domain reduced the interaction signal and the deletion of the NBD domain increased the interaction signal (Figure 1F and 1G). These results taken together indicate that pathologic α-Syn directly binds to hNOD2 primarily through the CARD domains.

Surface plasmon resonance (SPR), a technology that enables comprehensive and quantitative studies of protein-protein interactions (Hanson and Whelan, 2023), was used to determine the equilibrium binding kinetics between NOD2 and pathologic α-syn (Figure 1H). To ascertain the binding affinity, the Fc-tag from the tagged hNOD2 was removed using HRV3C protease (Figure S3A and S3B). Blue native page followed by α-syn immunoblotting confirmed the presence of aggregated α-syn and α-syn monomers (Figure S3C). α-Syn monomer or α-Syn PFFs (aggregates) as the ligand were immobilized on the sensor chip surface (Figure S3D and S3E), respectively. Monitoring the ligand (α-syn monomer or α-Syn PFF) – analyte (FL hNOD2) interaction in real-time and determining the association and dissociation constants allowed a determination of the Kd values (Figure 1I and 1J). The sensorgrams revealed an increase in the SPR signal (RU), reaching a steady state before the end of the injection of FL hNOD2, followed by rapid dissociation with the signal decreasing to zero during the successive injection of increasing concentrations, indicating high affinity binding (KD=8.07e-7M) between NOD2 and α- Syn PFF (Figure 1I). No significant binding affinity was observed with immobilized α-syn monomer (Figure 1J).

To determine whether FL hNOD2 binds to human pathologic α-Syn, human PD patient-derived pathologic α-Syn was generated by protein misfolding cyclic amplification (PMCA) technology. PMCA has been used to characterize pathologic α-Syn from human cerebrospinal fluid (CSF) (Kang et al., 2019; Shahnawaz et al., 2020; Shahnawaz et al., 2017). PMCA was performed for 7 days using CSF samples from both healthy controls and PD patients (Figure S3F and Table S2). α-Syn immunoblot analysis demonstrated increased high molecular weight α-Syn species in the PMCA products derived from PD patients compared with controls (Figure S3G and Table S2). To further characterize the PMCA-derived α-Syn species, transmission electron microscopy (TEM) and thioflavin T (ThT) fluorescence signals were assessed. The PD patient PMCA-derived α-Syn species displayed faster aggregation kinetics compared to controls (Figure S2H and S2I and Table S2). TEM images of PMCA products from the control group showed round-shaped aggregates with only a few rod-shaped aggregates, while the PD group primarily exhibited elongated rod-like fibrils, which were longer than those observed in controls (Figure S2J and S2K and Table S2). Immobilization of the α-Syn PMCA products from PD patients or healthy controls on a sensor chip surface was performed to monitor hNOD2 interactions (Figure 1H and Table S2). α-Syn PMCA products from PD patients displayed substantially higher affinity (KD=1.58e-6M) for FL hNOD2 compared to those from healthy controls (KD=4.56e-6M) (Figure 1K and 1L and Table S2). Together these results provide evidence that pathologic α-Syn binds to hNOD2 and monomeric α-Syn exhibits minimal binding to hNOD2. Moreover, pathologic α- Syn derived from PD patients binds to hNOD2.

### α-Syn PFF binding to NOD2 leads to recruitment and activation of RIPK2

NOD2 oligomerization through its NOD and CARD domains leads to the recruitment of RIPK2 through homotypic CARD-CARD interactions (Fridh and Rittinger, 2012; Ogura et al., 2001), generating an oligomeric signalosome, which is crucial for downstream signaling (Gong et al., 2018; Pellegrini et al., 2018). We wondered whether pathologic α-Syn binding to NOD2 (Figure 1F, 1G, S2E, and S2F) and its rapid dissociation (Figure 1I) could lead to its oligomerization, which in turn recruits RIPK2 through CARD-CARD interactions to initiate downstream signaling. Accordingly, the interaction between NOD2 and α-syn PFF over time was monitored (Figure S4A). BV2 microglia-like cells transfected with His- and Myc-tagged FL hNOD2 were treated with α- Syn-biotin PFF for 20 min, 1h, 3h, 6h, and 12h, followed by lysis and reciprocal pull-down using a NOD2 antibody and neutravidin, respectively (Figure S4A). Both pull-down analyses revealed that α-Syn PFF transiently binds to hNOD2 at 20 min and 1h, after which it dissociates from NOD2 at 3h (Figure S4B-S4E).

To determine whether α-Syn aggregates binding to NOD2 leads to complex with RIPK2, BV2 cells expressing both EGFP-fused hNOD2 (hNOD2-EGFP) and DSRED2-fused hRIPK2 (DSRED2-hRIPK2), as well as cells expressing both EGFP-fused hNOD1 (hNOD2-EGFP) and DSRED2-hRIPK2 as control were treated with α-syn PFF for 1h, 3h, and 6h followed by immunofluorescence detection (Figure S4F). Both hNOD2-EGFP and DSRED2-hRIPK2 proteins showed equal cytoplasmic localization and formed significantly larger complexes and a significant increase in the colocalization coefficient and area following α-Syn PFF treatment (Figure S4G-S4I). In contrast, BV2 cells expressing both hNOD1-EGFP and DSRED2-hRIPK2 showed no major difference in complex formation before and after α-Syn PFF treatment (Figure S4G-S4I).

To investigate whether α-Syn aggregates induce the formation of large NOD2/RIPK2 complexes, primary microglia were transduced with both lentivirus expressing HA-tagged hNOD2 (hNOD2-HA) and adeno-associated virus (AAV) expressing hRIPK2, followed by treatment with α-Syn PFF. The microglia were lysed using a non-ionic (NP-40) buffer and analyzed through a blue native polyacrylamide electrophoresis (BN-PAGE) assay (Figure S4J). Immunoblot analysis showed that hNOD2/hRIPK2 proteins formed large oligomeric complexes and displayed an increase in pS176 RIPK2 immunoreactivity, a marker of RIPK2 activation (Dorsch et al., 2006; Nachbur et al., 2015), after α-Syn PFF treatment (Figure S4K and S4L). Collectively, these findings indicate that pathologic α-Syn aggregates act as a ligand for NOD2 leading to the activation of the NOD2/RIPK2 signaling axis.

### α-Syn PFF activates NF-kB and MAPK through NOD2/RIPK2

Previous studies have shown that NOD2/RIPK2 activation leads to NF-κB (Nuclear Factor kappa B) transcriptional signaling (Ogura et al., 2001). Cell fractionation of wild type (WT), RIPK2^-/-^ and NOD2^-/-^ primary microglia activated with α-Syn PFF was separated into cytoplasmic and nuclear fractions to monitor RIPK2 and NF-κB activation (Figure 2A). In α-Syn PFF-treated WT microglia, pS176 RIPK2 immunoreactivity increased over time, peaking at 3 hours (Figure 2B and 2C, left panel). Additionally, α-Syn PFF induced the activation of the IkB kinase (IKK) complex and the degradation of IkBα (Figure 2B and 2C, left panel). The maximum nuclear translocation of p65 occurred when pS176 RIPK2 immunoreactivity was at the highest levels 3h after α-Syn PFF treatment (Figure 2B and 2C, left panel). Moreover, the total expression of RIPK2 protein significantly increased at 6h and 12h after α-Syn PFF treatment, and RIPK2 activation was sustained (Figure 2B and 2C, left panel). α-Syn PFF-treated RIPK2^-/-^ microglia showed significantly decreased NF-kB activation (Figure 2B and 2C, middle panel). In NOD2^-/-^ primary microglia total RIPK2 and pS176 RIPK2 immunoreactivity was significantly increased at 6h and 12h after α-Syn PFF treatment (Figure 2B and 2C, right panel). pS176 RIPK2 immunoreactivity was markedly reduced in NOD2^-/-^ microglia compared to WT microglia indicating that NOD2 is required for the full activation of RIPK2. Consistent with this, α-Syn PFF-induced NF-κB signaling activation was partially blocked in NOD2^-/-^ microglia (Figure 2B and 2C, right panel).

**Figure 2.**
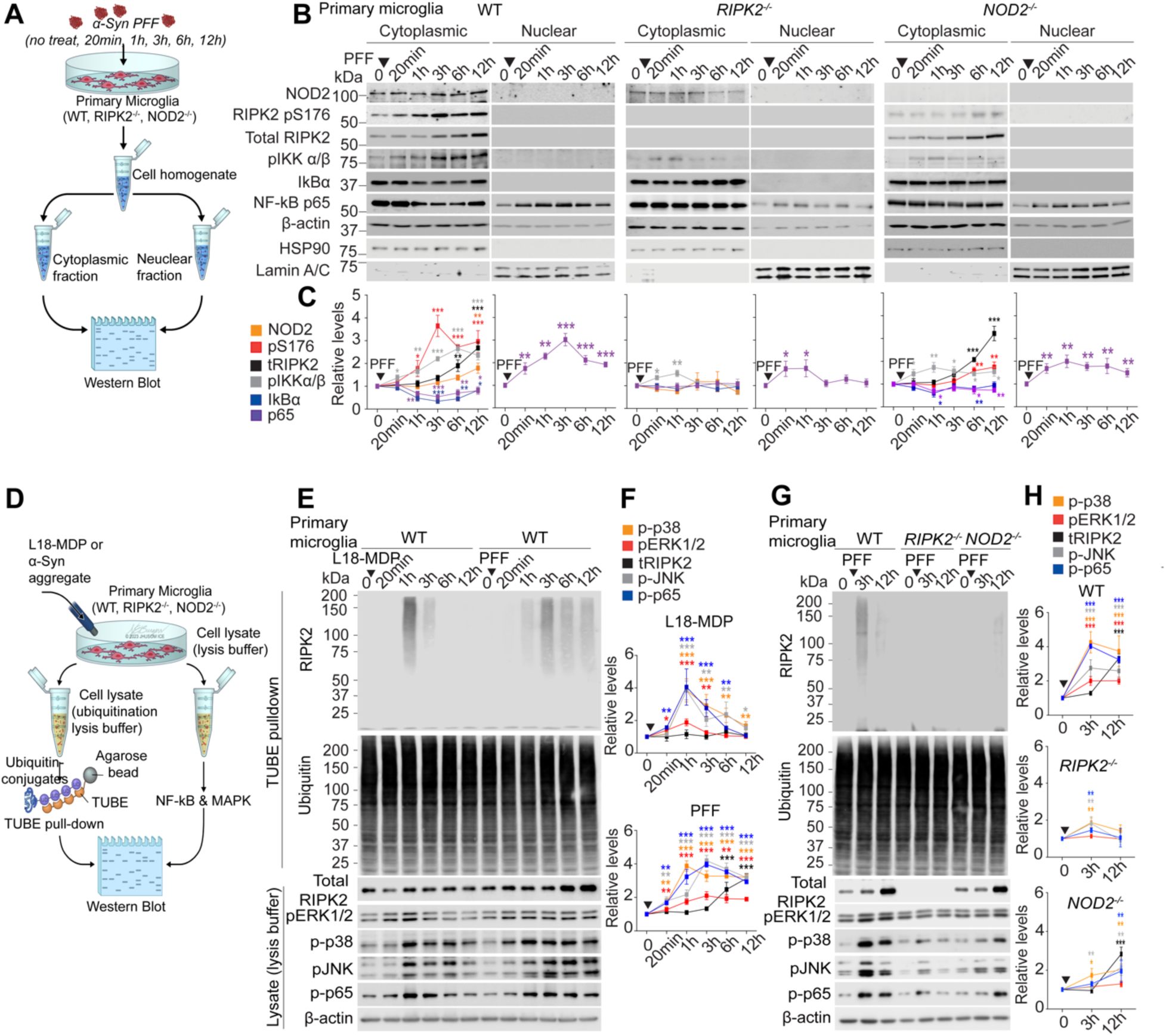
α-Syn PFF enhances NF-κB and MAPK activation through RIPK2 ubiquitination. (A) Schematic diagram showing immunoblotting with specific antibodies to examine NF-κB activation and nuclear translocation after cytoplasmic and nuclear fractionation of WT, RIPK2^-/-^, or NOD2^-/-^ primary microglia treated with α-Syn PFF (5 μg/ml) for 20 min, 1h, 3h, 6h, or 12h. (B) Immunoblots revealed significant inhibition of α-Syn PFF-induced NF-kB activation in the absence of RIPK2 or NOD2. (C) Protein level quantification in B shown as a line graph (n=3, biologically independent experiments), with data presented as mean ± SEM (*P < 0.05, **P < 0.01, ***P < 0.001). (D) Schematic diagram showing TUBE pull-down and MAPK and NF-κB immunoblot analysis to confirm ubiquitinated RIPK2 and downstream signaling activation in cell lysates of *WT, RIPK2^-/-^*, or *NOD2^-/-^* primary microglia treated with either M18-MDP (1 μg/ml) or α-Syn PFF (5 μg/ml) for 20 min, 1h, 3h, 6h, or 12h. (E) Representative immunoblots of RIPK2, ubiquitin, MAPK signaling, and phospho-p65 after M18-MDP or α-Syn PFF treatment. (F) Quantification of protein levels in (E) presented as a line graph (n=3, biologically independent experiments). Data shown as mean ± SEM (*P < 0.05, **P < 0.01, ***P < 0.001). (G) Immunoblots of α-Syn PFF-induced ubiquitination of RIPK2 and downstream signaling in RIPK2^-/-^ or NOD2^-/-^ primary microglia. (H) Quantification of protein levels in (G) shown as a line graph (n=3, biologically independent experiments). Data shown as mean ± SEM (*\*P* < 0.05, *\*\*P* < 0.01, *\*\*\*P* < 0.001).

NOD2 stimulation leads to robust ubiquitination of RIPK2, which is critical for the formation of the signaling complex and effective downstream pathway activation (Damgaard et al., 2012; Yang et al., 2007). To determine whether pathologic α-Syn drives RIPK2 ubiquitination, a tandem ubiquitin binding entity (TUBE) pull-down assay was used to purify ubiquitinated proteins from primary microglia stimulated with the NOD2 ligand L18-MDP or α-Syn PFF. This assay allows for the efficient detection of polyubiquitylated proteins in their native state (Hjerpe et al., 2009; Lopitz-Otsoa et al., 2012) and facilitates the monitoring of activation of downstream signaling events (Figure 2D). L18-MDP-treated WT primary microglia exhibited rapid ubiquitination of RIPK2, which returned to baseline levels after 6 hours (Figure 2E). Additionally, L18-MDP induced MAPK ERK/p38/JNK and NF-κB p65 activation (Figure 2E and 2F). α-Syn PFF-activated WT primary microglia exhibited rapid and sustained RIPK2 ubiquitination, although the onset was delayed compared with L18-MDP treatment (Figure 2E and 2F). While L18-MDP-induced MAPK and NF-κB signaling activation returned to basal level in a time-dependent manner, α-Syn PFF-treated WT microglia showed sustained MAPK and NF-κB activation (Figure 2E and 2F). α-Syn PFF-treated RIPK2^-/-^ microglia showed a complete inhibition of RIPK2 ubiquitination confirming the specificity of the assay and a significant blockade of MAPK and NF-κB activation (Figure 2G and 2H). In NOD2^-/-^ primary microglia, α-Syn PFF induced the ubiquitination of RIPK2 was markedly reduced (Figure 2G and 2H). Collectively, these results demonstrate that α-Syn PFF induces acute activation of RIPK2 through NOD2 stimulation and activation of NF-κB.

### NOD2/RIPK2 signaling is activated in PD

The levels of *NOD2 and RIPK2 mRNA* and protein as well as RIPK2 activation (pS176 RIPK2) were assessed in the substantia nigra (SN) of PD patient versus control postmortem brain (Table S2). Microglia reactivity as assessed by Iba immunoreactivity was observed in PD substantia nigra along with increased pS129 α-Syn immunoreactivity and reduced TH immunoreactivity (Figure 3A-3D). Both NOD2 and RIPK2 mRNA and protein levels were significantly elevated in the SN of PD patients compared to healthy controls (Figure 3E-3G and Table S2). Furthermore, western blot analysis revealed an increase in pS176 RIPK2 immunoreactivity in the SN of PD patients (Figure 3F and 3G and Table S2). PLA using specific antibodies for pathologic α-Syn and NOD2 protein as described in Figure 1A was used to explore the interaction between pathologic α-Syn aggregates and NOD2 in microglia in the SN of PD versus control patient tissue slices (Figure 3H and Table S2). Consistent with our previous PLA results shown in α-Syn PFF-treated primary microglia (Figure 1A-1C), we observed strong fluorescence signals in Iba1-positive microglia in the SN of PD patients, whereas these signals were absent in healthy controls (Figure 3I and 3J and Table S2). Immunofluorescence analysis revealed that pS176 RIPK2 immunoreactivity was significantly colocalized in Iba-1-positive microglia (Figure 3K and 3N and 3O and Table S2) compared to GFAP (Figure 3L and 3N and Table S2) and TH immunoreactivity (Figure 3M and 3N and Table S2) in the SN of PD and healthy control patients, while there is significantly more colocalization with GFAP compared to TH immunoreactivity (Figure 3N). Within Iba-1-positive microglia, pS176 RIPK2 immunoreactivity was significantly increased in the SN of PD compared to healthy control patients (Figure 3O and Table S2).

**Figure 3.**
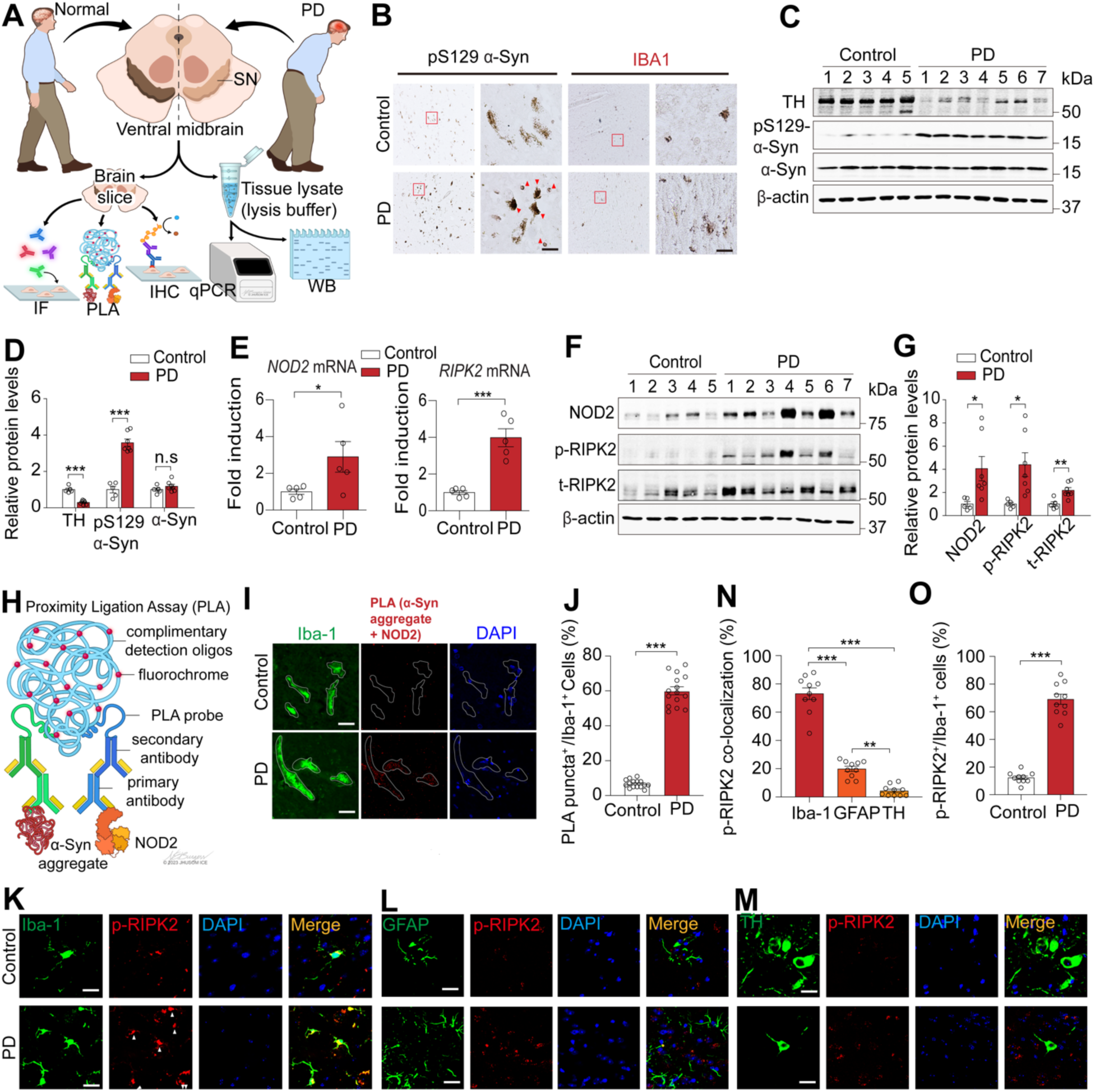
NOD2/RIPK2 signaling is activated in PD patients. (A) Illustration depicting biochemical analyses to identify NOD2/RIPK2 signaling activation in postmortem brain tissues of PD patients and healthy controls. (B) Representative photomicrographs showing Lewy bodies in pigmented neurons (left images) and reactive microglia (right images) in the substantia nigra (SN) of PD versus control patients. Red triangles indicate the Lewy bodies. (C) Immunoblots showing reduced TH protein and elevated pS129 α-syn levels in the substantia nigra (SN) of human PD versus control postmortem brain. (D) Bar graph quantifying protein expression in C (n=5 for healthy control, n=7 for PD patient). Data are presented as mean ± SEM (***P < 0.001). (E) Bar graph quantifying *NOD2* and *RIPK2* mRNA levels in the substantia nigra (SN) of PD patients versus controls using qPCR (n=5 in each group). Data are presented as mean ± SEM (*P < 0.05, ***P < 0.001). (F) Immunoblots showing the accumulation of NOD2, total RIPK2 (t-RIPK2), and elevated pS176 RIPK2 levels in PD versus control postmortem brain. (G) Bar graph quantifying protein expression in F (n=5 for healthy control, n=7 for PD patient). Data are presented as mean ± SEM (*P < 0.05). (H) Schematic diagram depicting the PLA to validate the interaction between endogenous pathologic α-syn aggregates and NOD2 protein in microglia of PD postmortem brain. (I) Confocal images displaying the detection of PLA-positive signals (red dots) in the postmortem brain slices of PD patients, but not in those of healthy controls. (J) Bar graph quantifying the percentage of PLA-positive/total Iba1-positive cells in I (n=15 in each group). Data are presented as mean ± SEM (***P < 0.001). (K) Confocal images showing significant co-localization of pS176 RIPK2 with Iba-1 signals in PD versus control patients. White triangles indicate the accumulation of p-RIPK2 in microglia. (L and M) Confocal images demonstrating minimal co-localization of pS176 RIPK2 with GFAP and TH signals. (N) Bar graph quantifying the percentage of co-localization in K-M. Data are presented as mean ± SEM (*\*\*P* < 0.01, *\*\*\*P* < 0.001). (O) Bar graph quantifying the percentage of p-RIPK2 positive/total Iba-1 positive cells in K. Data are presented as mean ± SEM (*\*\*\*P* < 0.001).

### NOD2 or RIPK2 deletion prevents NF-κB and MAPK activation in the α-Syn PFF mouse model of PD

The role of NOD2 and RIPK2 on NF-κB and MAPK activation was assessed in intrastriatal α-Syn PFF injected mice, a model of sporadic PD (Luk et al., 2012a; Luk et al., 2012b; Mao et al., 2016). α-Syn PFF was stereotaxically injected into the striatum of WT, RIPK2^-/-^, and NOD2^-/-^ mice (Figure 4A). Six months post-injection, immunoblot analysis revealed that NOD2 immunoreactivity was significantly elevated in the ventral midbrain of α-Syn PFF-injected WT, but not RIPK2^-/-^ and NOD2^-/-^ mice (Figure 4B and 4C). RIPK2 and pS176 RIPK2 were significantly elevated in the ventral midbrain of α-Syn PFF-injected WT, but not RIPK2^-/-^ mice (Figure 4B and 4C). Although RIPK2 and pS176 RIPK2 were significantly elevated in the ventral midbrain of α- Syn PFF-injected NOD2^-/-^ mice, the level of elevation was substantially less than that of WT mice (Figure 4B and 4C). Immunofluorescence analysis revealed a significant increase in pS176 RIPK2 in Iba-1 positive cells in the ventral midbrain that was substantially greater than the significant increase in pS176 RIPK2 in GFAP positive cells in α-syn PFF-injected WT mice (Figure 4D and 4E). The increase in pS176 RIPK2 immunoreactivity in Iba-1 positive cells was significantly reduced in RIPK2^-/-^ and NOD2^-/-^ mice (Figure 4D and 4E). The increase in pS176 RIPK2 immunoreactivity in GFAP positive cells was eliminated in RIPK2^-/-^ mice and reduced in NOD2^-/-^ mice (Figure 4D and 4E), consistent with the human results (Figure 3). No detectable pS176 RIPK2 immunoreactivity was observed in TH-positive neurons (Figure 4D and 4E). To confirm that α-Syn PFF primarily activates NOD2 and RIPK2 signaling in microglia, we monitored RIPK2 and pS176 RIPK2 immunoreactivity via immunoblot analysis in mouse primary microglia, astrocytes, and neurons treated with α-Syn PFF (Figure S5A). Consistent with our human and *in vivo* mouse histological findings (Figure 3 and Figure 4D and 4E), pS176 RIPK2 immunoreactivity was predominantly increased in α-Syn PFF-treated primary microglia (Figure S5B and S5C). NOD2/RIPK2 proteins were primarily expressed in primary microglia, with minimal detection in primary astrocytes (Figure S5B and S5C). NOD2, RIPK2, and pS176 RIPK2 immunoreactivity were barely detectable in the primary neurons (Figure S5B and S5C). PLA using specific antibodies for pathologic α-Syn and NOD2 protein as described in Figure 1A also revealed that the binding between NOD2 and α-Syn aggregates predominantly occurs within microglial cells (Figure S5D-S5F).

**Figure 4.**
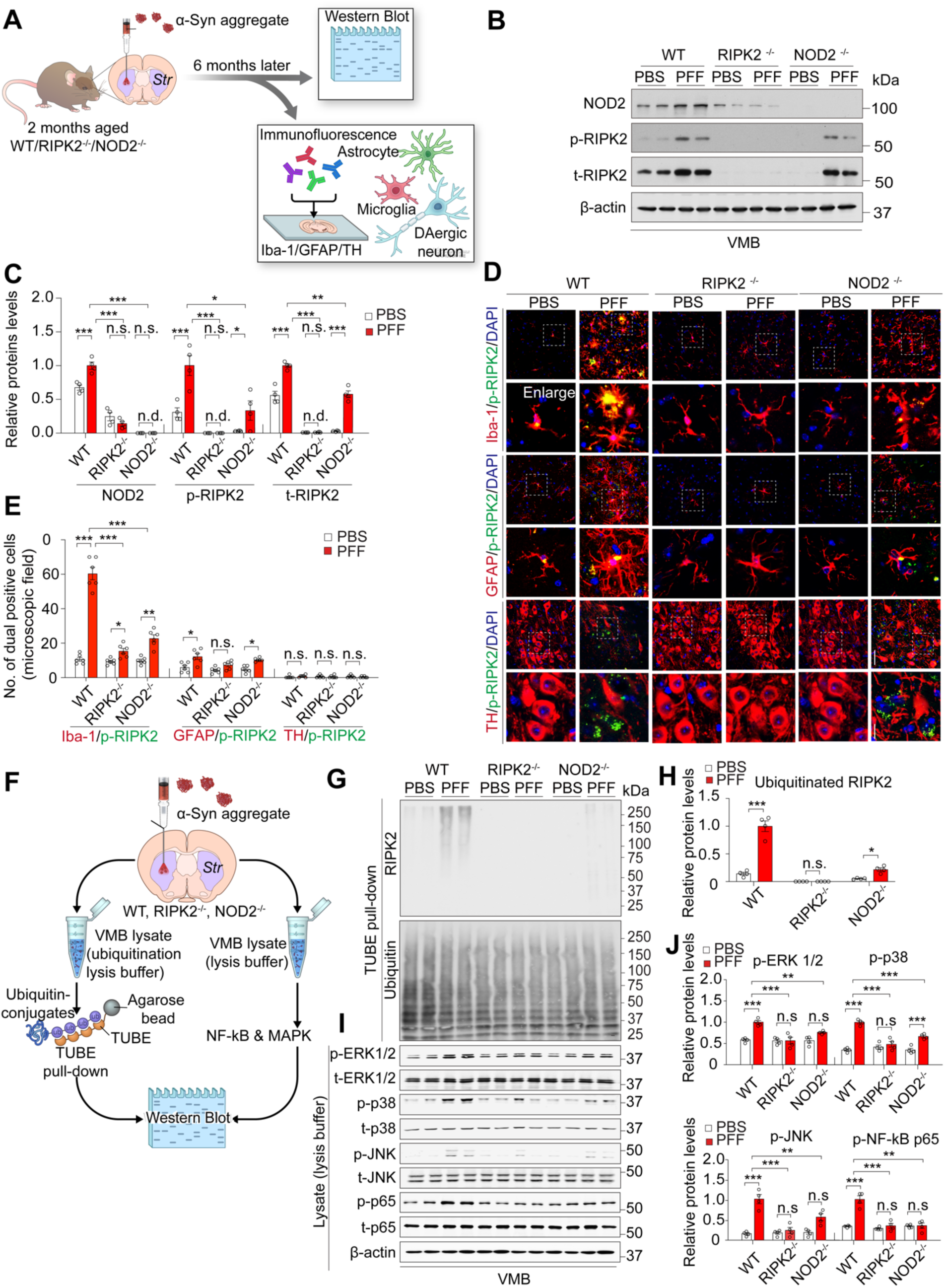
Deletion of NOD2 or RIPK2 in α-Syn PFF-injected PD mice reduces microglial signaling and NF-κB and MAPK by blocking RIPK2 ubiquitination. (A) Schematic diagram depicting the biochemical analyses to study NOD2 and RIPK2 signaling activation and neuroinflammation 6 months after α-Syn PFF injection in the ventral midbrain of WT, RIPK2^-/-^, and NOD2^-/-^ mice. (B) Representative immunoblots of NOD2, t-RIPK2, and p-RIPK2 in α-Syn PFF-injected WT, RIPK2^-/-^, and NOD2^-/-^ mice. (C) Bar graph quantifying protein levels in B (n=4 in each group). Data are presented as mean ± SEM (n.s; not significant, *\*P < 0.05, **P < 0.01, ***P < 0.001*). (D) Confocal images of assessment of co-localization of pS176 RIPK2 with Iba-1, GFAP, and TH in α-Syn PFF-injected WT, RIPK2^-/-^ and NOD2^-/-^ mice. Specific areas are enlarged (dotted line). (E) Bar graph quantifying the number of double-positive cells of Iba-1, GFAP, or TH with p-RIPK2 in D (n = 6 in each group). Data are presented as mean ± SEM (n.s; not significant, *\*P < 0.05, **P < 0.01, ***P < 0.001*). (F) Schematic diagram showing TUBE pull-down and MAPK and NF-κB western blot analysis to detect ubiquitinated RIPK2 and downstream signaling activation in the ventral midbrain of α-Syn PFF-injected WT, RIPK2^-/-^, or NOD2^-/-^ mice. (G) Representative immunoblots depicting α-Syn PFF-induced RIPK2 ubiquitination in WT, RIPK2^-/-^ or NOD2^-/-^ mice. (H) Quantification of ubiquitinated RIPK2 protein levels in G presented as a bar graph (n=4, per group). Data are expressed as mean ± SEM (*P < 0.05, ***P < 0.001). (I) Immunoblots of α-Syn PFF-induced MAPK and NF-κB activation in *WT, RIPK2^-/-^ or NOD2^-/-^* mice. (J) Quantification of protein levels in I (VMB lysate using lysis buffer) presented as a bar graph (n=4, per group). Data are expressed as mean ± SEM (**P < 0.01, ***P < 0.001).

The impact of NOD2 or RIPK2 depletion on α-Syn PFF-induced microglial NOD2 and RIPK2 signaling and downstream activation was investigated in α-Syn PFF-injected mice (Figure 4F). A significant increase in RIPK2 ubiquitination accompanied by MAPK and NF-kB activation was observed in WT mice with a considerable reduction in MAPK and NF-kB activation and blockade of ubiquitination of RIPK2 in RIPK2^-/-^ and NOD2^-/-^ mice (Figure 4G-J). These findings strongly support that pathologic α-Syn induced NOD2 and RIPK2 activation is enriched in microglia and that downstream pathway activation is effectively suppressed by the genetic depletion of NOD2 and RIPK2.

### Deletion of NOD2 or RIPK2 prevents neuroinflammation

To investigate whether genetic deletion of NOD2 or RIPK2 prevents glial activation induced by α-Syn PFF, microglia, and astrocyte activation were assessed using immunohistochemistry (IHC) for Iba-1 and GFAP respectively. α-Syn PFF intrastriatal injected WT mice showed a significant increase in Iba-1 and GFAP immunoreactivity 6 months post injection, consistent with previous findings (Yun et al., 2018). The increase in Iba-1 and GFAP immunoreactivity was effectively blocked in RIPK2^-/-^ and NOD2^-/-^ mice (Figure S5G and S5H). Moreover, immunoblot analysis confirmed that the protein levels of Iba-1 and GFAP were significantly increased in the ventral midbrain of α-Syn PFF-injected WT mice while the elevation in Iba-1 was eliminated in RIPK2^-/-^ mice and substantially reduced in NOD2^-/-^ mice (Figure S5I and S5J). The elevation in GFAP was completely attenuated in both RIPK2^-/-^ and NOD2^-/-^ mice (Figure S5I and S5J). These results indicate that NOD2 and RIPK2 play a critical role in α-Syn PFF-induced glial activation, and their genetic deletion can effectively mitigate the neuroinflammatory response.

### Deletion of NOD2 or RIPK2 prevents α-Syn PFF induced neurotoxic reactive astrocytes

Microglia induce neurotoxic reactive astrocytes by releasing proinflammatory factors such as interleukin 1α (IL-1α), tumor necrosis factor (TNF), and complement component 1, subcomponent q (C1q) (Guttenplan et al., 2021; Liddelow et al., 2017) and α-Syn PFF-induced neuroinflammatory microglia can convert naive astrocytes to neurotoxic reactive astrocytes by secreting these cytokines (Yun et al., 2018). To determine whether microglial NOD2 and RIPK2 signaling contribute to the formation of neurotoxic reactive astrocytes induced by pathologic α- Syn PFF, C3 immunoreactivity of GFAP-positive astrocytes, which is characteristic of neurotoxic reactive astrocytes was monitored (Liddelow et al., 2017) (Figure 5A). Conditioned medium from microglia exposed to α-Syn PFF (PFF-MCM) significantly increased the levels of GFAP and C3, while conditioned medium from microglia treated with PBS (PBS-MCM) had no effect (Figure 5B, 5C, S6A and S6B). This effect was effectively blocked by the absence of RIPK2 and substantially reduced by the absence of NOD2 in microglia (Figure 5B, 5C, S6A, and S6B). The effect of deletion of NOD2 or RIPK2 in microglia on the secretion of factors that induce neurotoxic reactive astrocytes was monitored via quantitative polymerase chain reaction (qPCR) (Figure 5A). Primary microglia from WT, NOD2^-/-^ or RIPK2^-/-^ mice were treated with α-Syn PFF and mRNA levels of *TNFα, IL-1α, IL-1β, C1q,* and *IL-6* were measured using qPCR (Figure 5A). α-Syn PFF-treated *WT* microglia showed significant increases in mRNA levels, for *C1q, TNFα, IL-1α, IL-1β*, and *IL-6*, which were markedly inhibited by the absence of NOD2 or RIPK2 (Figure 5D). The effects of the deletion of NOD2 or RIPK2 in microglia on the expression of neurotoxic reactive transcripts in astrocytes, which are upregulated by reactive microglia (Anderson et al., 2016; Liddelow et al., 2017; Zamanian et al., 2012) was monitored. PBS- or PFF-MCM from WT, NOD2^-/-^ or RIPK2^-/-^ microglia was applied to primary WT astrocytes, and pan-, A1-like and A2-like mRNA levels were measured using qPCR (Figure 5A). As previously described (Yun et al., 2018), PFF-MCM from WT microglia significantly increased pan- and A1-like transcripts in primary astrocytes, which was attenuated by the absence of NOD2 or RIPK2 in microglia (Figure 5E). To investigate whether deletion of NOD2 or RIPK2 could inhibit the formation of these astrocytes induced by α-Syn PFF *in vivo* (Figure 5F), C3 or GFAP immunoreactivity in the substantia nigra (SN) of α-Syn PFF-injected mice was assessed. α-Syn PFF-injected WT mice showed a significant increase in C3 or GFAP immunoreactivity, which was reduced in NOD2^-/-^ mice and eliminated in RIPK2^-/-^ mice (Figure 5G and 5H). Moreover, the formation of C3- and GFAP-dual positive astrocytes in the SN of α-Syn PFF-injected WT mice as determined by immunofluorescence was reduced in NOD2^-/-^ mice and eliminated in RIPK2^-/-^ mice (Figure S6C and S6D). Microglia and astrocytes were enriched by immunopanning from the ventral midbrain of α-Syn PFF-injected WT, NOD2^-/-^ or RIPK2^-/-^ mice, and qPCR was performed to assess the levels of *C1q, TNFα, IL-1α, IL-1β*, and *IL-6* in microglia (Figure 5I) and pan-, A1-like and A2-like mRNA levels in astrocytes, as described previously (Yun et al., 2018). The deletion of NOD*2* or *RIPK2* significantly inhibited the expression of *C1q, TNFα, IL-1α, IL-1β*, and *IL-6* in microglia (Figure 5I) and neurotoxic reactive transcripts in astrocytes (Figure 5J) induced by α-Syn PFF *in vivo*.

**Figure 5.**
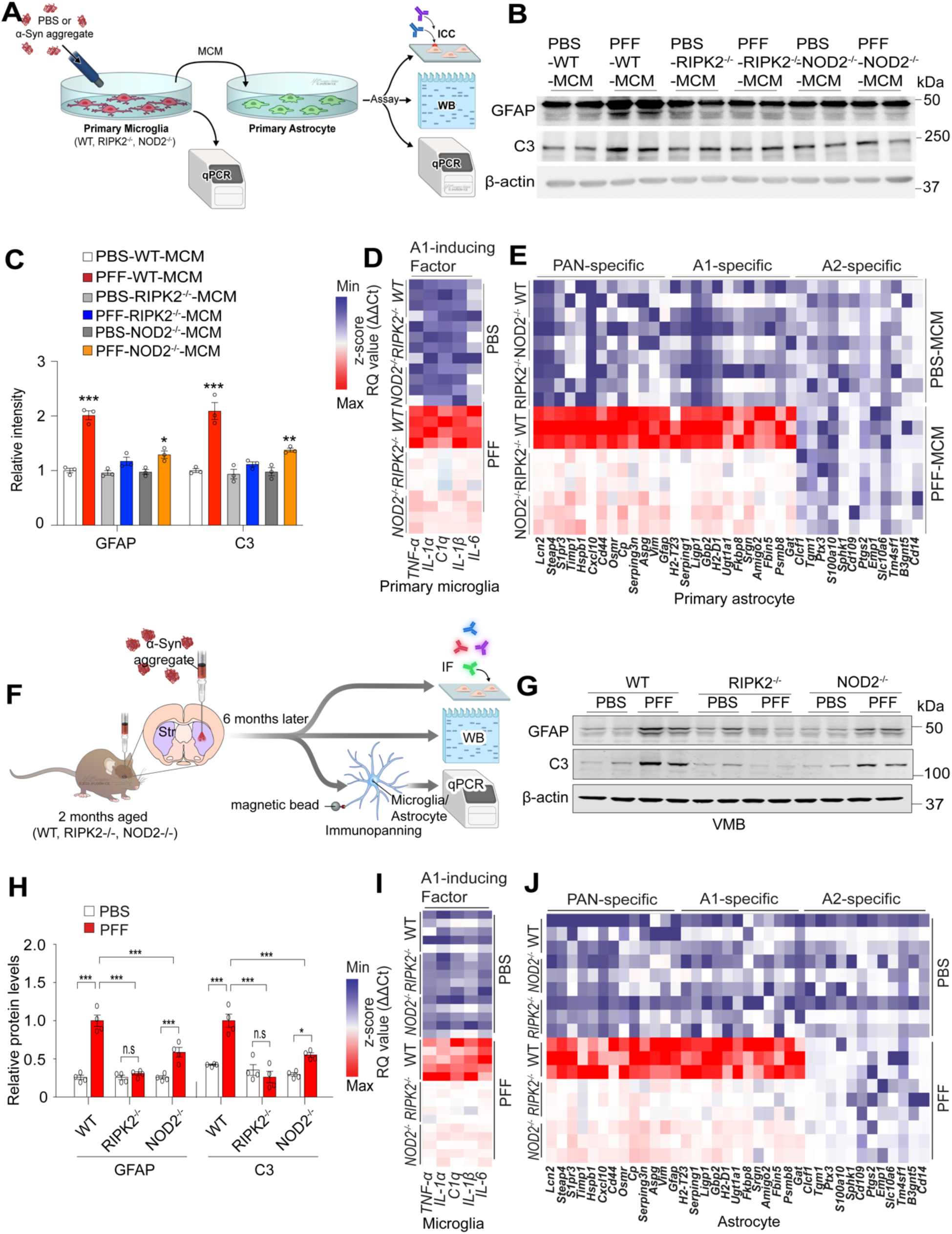
NOD2 or RIPK2 deletion suppresses α-Syn PFF-induced microglial activation and neurotoxic reactive astrocyte formation. (A) Schematic diagram illustrating the experimental processes in astrocytes and microglia after treatment with MCM from α-Syn PFF-treated WT, NOD2^-/-^, and RIPK2^-/-^ primary microglia. (B) Representative immunoblots of GFAP and C3 protein levels in astrocytes due to PFF-MCM in WT, NOD2^-/-^ or /RIPK2^-/-^ microglia. (C) Bar graph representing the quantification of protein levels in (B) (n=3, independent experiments). Data are presented as mean ± SEM (*P < 0.05, **P < 0.01, ***P < 0.001). (D) Heatmap representing mRNA transcripts level of C1q and proinflammatory cytokine genes using qPCR in WT, NOD2^-/-^, and RIPK2^-/-^ primary microglia treated with PBS or α-Syn PFF. (E) Heatmap showing PAN, A1, and A2-specific transcript levels of primary astrocytes incubated with MCM collected from WT, NOD2^-/-^, and RIPK2^-/-^ primary microglia treated with PBS or α-Syn PFF. (F) Illustration depicting the experimental design to investigate α-Syn PFF-induced microglia activation and neurotoxic reactive astrocyte in the ventral midbrain of WT, NOD2^-/-^, and RIPK2^-/-^ mice. (G) Representative immunoblots of α-Syn PFF-induced increase in GFAP and C3 protein levels in WT, NOD2^-/-^, and RIPK2^-/-^ mice. (H) Bar graph presenting the quantification of protein levels in (G) (n=4, each group). Data are represented as mean ± SEM (*P < 0.05, ***P < 0.001). (I) Heatmap representing the reactivity of neurotoxic reactive astrocyte inducers from microglia in α-Syn PFF-injected WT, NOD2^-/-^ and RIPK2^-/-^ mice. (J) Heatmap from WT, NOD2^-/-^ and RIPK2^-/-^ mice of PAN-specific, A1-specific, and A2-specific transcripts following α-Syn PFF injection.

Neurotoxic reactive astrocytes are linked to detrimental effects on neuronal survival, outgrowth, synaptogenesis, and increased neuronal cell death (Liddelow et al., 2017). Previous research demonstrated that neurons co-cultured with neurotoxic reactive astrocytes exhibit reduced synapses and diminished synaptic activity compared to neurons co-cultured with naive astrocytes, indicating a negative impact on neuronal activity and connectivity (Liddelow et al., 2017). To explore whether α-Syn PFF-induced neurotoxic reactive astrocytes inhibit the development of spontaneous neuronal activity, a multi-electrode array (MEA) system was utilized to monitor the spontaneous firing of neurons in vitro (McConnell et al., 2012; Negri et al., 2020) (Figure 6A). The firing rate of neurons treated with astrocyte conditioned media (ACM) from PBS- or PFF-MCM treated astrocytes was analyzed on day in vitro (DIV) 7, 10, and 14 in which the neuronal cultures received ACM from astrocytes treated with PBS- or PFF-MCM and at DIV 7 and 10 (Figure 6B-6D). Neurons cultured with ACM prepared from WT microglia treated with PFF-MCM did not display a significant development of mean firing rate (Figure 6B and 6C) and burst frequency (Figure 6B and 6D) at DIV 10 and 14 when compared to neuronal cultures that received ACM from astrocytes treated with PBS-MCM. No significant impairment of mean firing rate (Figure 6B and 6C) and burst frequency (Figure 6B and 6D) at DIV 10 and 14 was observed in neurons treated with ACM from PBS- or PFF-MCM from NOD2^-/-^ or RIPK2^-/-^ microglia. Next, the ability of ACM from PBS- or PFF-MCM from WT, NOD2^-/-^ or RIPK2^-/-^ microglia to kill neurons was assessed (Figure 6A). As previously reported, ACM generated from WT PFF-MCM significantly increased neuronal cell death compared to WT PBS-MCM (Yun et al., 2018) as assessed by lactate dehydrogenase (LDH) release (Figure 6E) and via an Alamarblue assay (Figure 6F), while ACM prepared from PBS- or PFF-MCM from NOD2^-/-^ or RIPK2^-/-^ microglia failed to cause neuronal cell death (Figure 6E and 6F). Taken together these findings indicate a critical role for microglial NOD2 and RIPK2 in the formation of neurotoxic reactive astrocytes induced by α-Syn PFF-activated microglia.

**Figure 6.**
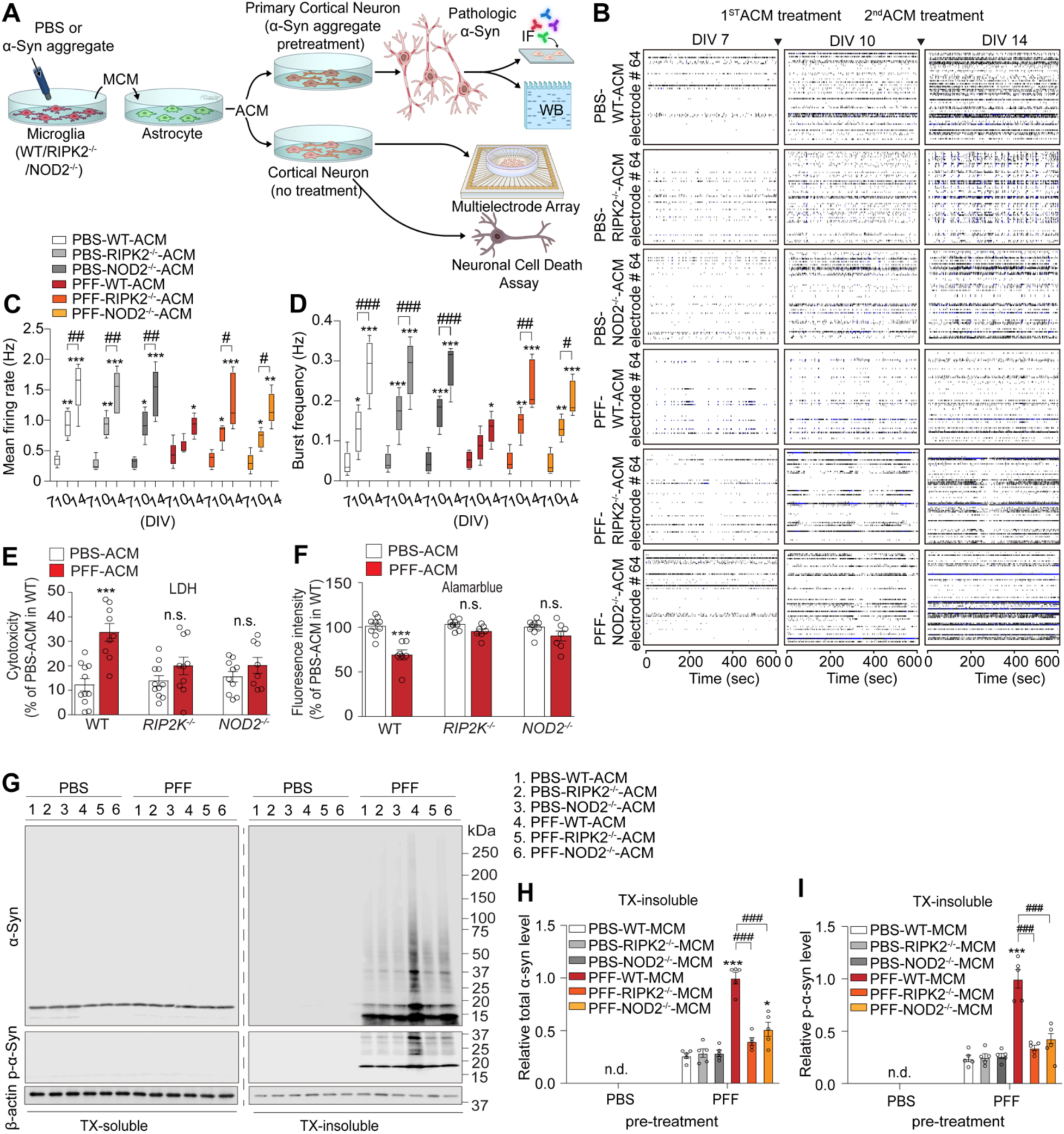
*NOD2/RIPK2 KO* protects against α-Syn PFF-induced neuropathology and neuronal loss. (A) Schematic diagram illustrating biochemical analyses for evaluating the impact of the deletion of NOD2 or RIPK2 on neuronal toxicity induced by PFF-ACM treatment into primary cortical neurons pre-treated with or without α-Syn PFF. (B) Representative raster plots showing the rescue effect of *RIPK2/NOD2 KO* on impaired electrophysiological maturation of neurons due to PFF-ACM. Dots represent peak electric signal positions over time from 64 MEA electrodes. (C and D) Quantification of mean firing rate (Hz) and burst frequency (Hz) in (B) shown as box plots. (n=5, biologically independent experiments). Data presented as median, interquartile range, minimum, and maximum range. (*P < 0.05, **P < 0.01, ***P < 0.001 - for comparison between DIV 7 and DIV 10; ^#^P < 0.05, ^##^P < 0.01, ^###^P < 0.001 - for comparison between DIV 10 and DIV 14). (E and F) Bar graph quantifying cytotoxicity levels and indicating the inhibitory effect of *RIPK2/NOD2 KO* on neuronal cell death due to neurotoxic astrocytes, as determined by LDH and Alamarblue assays (n=11, biologically independent experiments). Data presented as mean ± SEM (n.s; not significant, ***P < 0.001). (G) Representative immunoblots showing the inhibitory effect of *RIPK2/NOD2 KO* on the increase of Lewy-like pathology induced by PFF-ACM. (H and I) Bar graph quantifying total α-syn and p-α-syn protein levels in the Triton X-100-insoluble fraction in B (n=5, biologically independent experiments). Data presented as mean ± SEM (n.d. - no detection, *P < 0.001, ***P < 0.001, for comparison to α-Syn PFF-untreated group in microglia; ^###^P < 0.001, for comparison to α-Syn PFF-treated group in WT microglia).

### NOD2 or RIPK2 deletion prevents α-Syn PFF induced neurodegeneration

To determine whether neurotoxic reactive astrocytes might contribute to the formation of pathologic α-Syn, cortical neurons were treated with α-Syn PFF followed by treatment with ACM from PBS- or PFF-MCM from WT, NOD2^-/-^ or RIPK2^-/-^ microglia (Figure 6A). ACM from WT PFF-MCM significantly increased the accumulation of Triton X-100-insoluble α-Syn and pS129 α-Syn in WT cortical neurons treated with α-Syn PFF compared to ACM from WT PBS-MCM (Figure 6G-6I). ACM from PBS- or PFF-MCM from NOD2^-/-^ or RIPK2^-/-^ microglia failed to lead to a significant accumulation of Triton X-100-insoluble α-Syn and pS129 α-Syn in WT cortical neurons treated with α-syn PFF (Figure 6G-6I). Immunoreactivity for pS129 α-Syn in WT MAP2 cortical neurons confirmed that ACM from WT PFF-MCM significantly enhanced the immunoreactivity for pS129 α-Syn while ACM from PBS-WT MCM or PBS- and PFF-MCM from NOD2^-/-^ or RIPK2^-/-^ microglia failed to increase the immunoreactivity for pS129 α-syn in WT cortical neurons treated with α-syn PFF (S7A-S7C). These results taken together suggest that NOD2 and RIPK2 mediated activation of neurotoxic reactive astrocytes contribute to the formation of pathologic α-Syn.

The effects of NOD2 or RIPK2 deletion on pathologic α-Syn accumulation, dopaminergic (DA) neurodegeneration, and motor behavioral deficits were assessed in the intrastriatal α-Syn PFF model of PD (Figure 7A). 6 months after intrastriatal injection of α-Syn PFF, mice were evaluated for neuropathology, loss of DA neurons, and behavioral impairments (Figure 7A). Consistent with prior findings (Yun et al., 2018), histological studies revealed that α-Syn PFF-injected *WT* mice showed a substantial increase in pS129 α-Syn immunoreactivity in the ventral midbrain, indicating effective templating of endogenous α-Syn into pathologic α-Syn as assessed by pS129 α-Syn immunoreactivity (Figure 7B and 7C and Figure S7D and S7E). Immunoreactivity for pS129 α-Syn was significantly reduced in the ventral midbrain of α-Syn PFF-injected NOD2^-/-^ or RIPK2^-/-^ mice, particularly in tyrosine hydroxylase (TH) immunoreactive neurons (Figure S7D and S7E). Immunoblot analysis from Triton X-100-soluble and -insoluble fractions on the ventral midbrain of control or α-Syn PFF intrastriatal injected mice, revealed accumulation of Triton X- 100-insoluble α-Syn and pS129 α-Syn in the ventral midbrain of α-Syn PFF-injected WT mice (Figure 7D and 7E).

**Figure 7.**
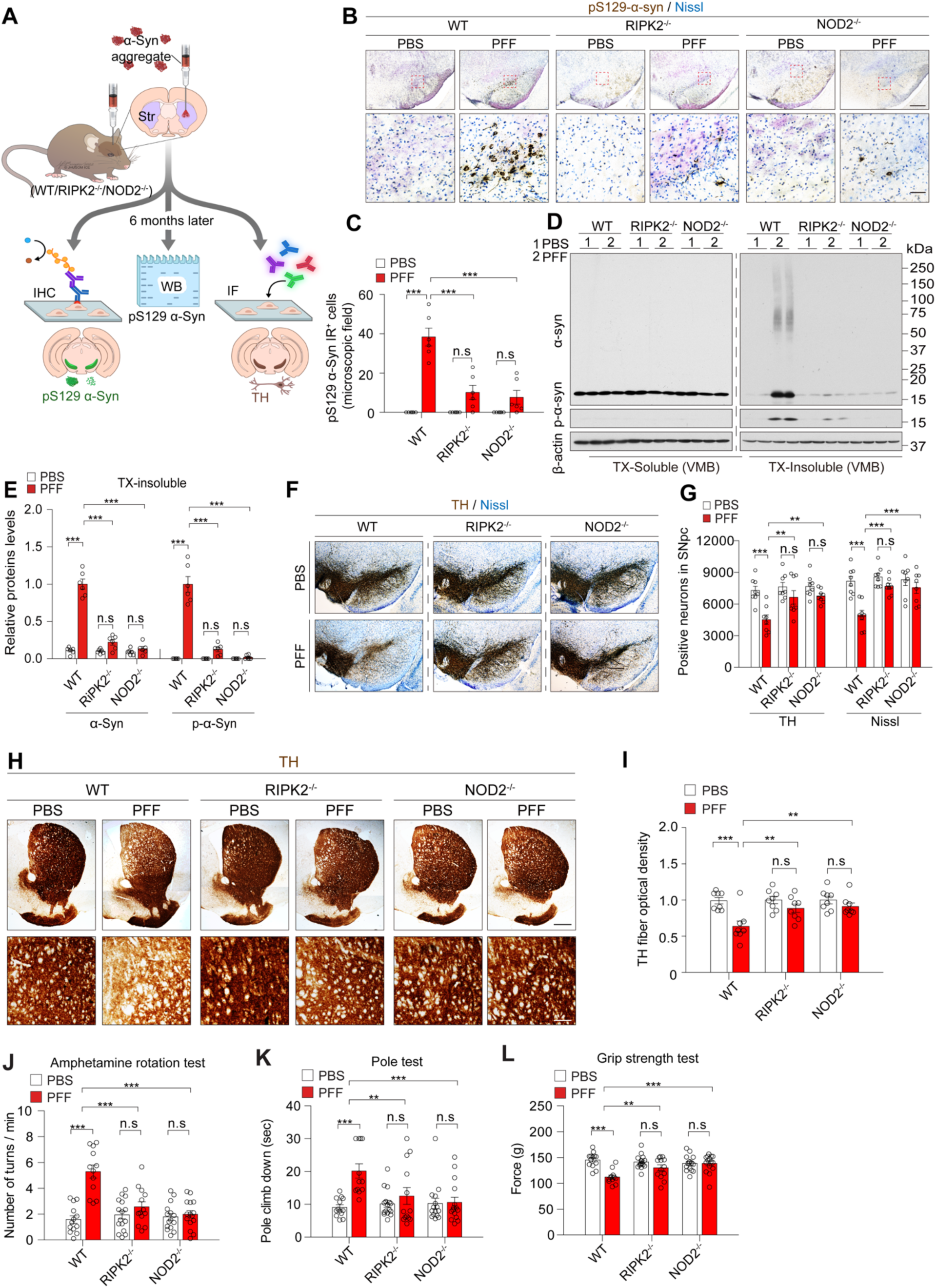
*NOD2/RIPK2 KO* protects against α-Syn PFF-induced neuropathology and dopaminergic neurodegeneration. (A) Schematic diagram depicting the evaluation of the inhibitory effect of *RIPK2/NOD2 KO* on Parkinson’s disease-like pathology in α-Syn PFF intra-striatal injected mice using biochemical study, immunofluorescence, and immunohistology with a stereotaxic instrument. (B) Representative immunohistochemistry images showing genetic depletion of RIPK2/NOD2 protects against α-Syn PFF-induced LB-like pathology (C) Bar graph displaying the number of p-α-Syn positive dopaminergic neurons in (B) (n = 6, each group). Data presented as mean ± SEM (n.s - not significant, ***P < 0.001). (D) Representative immunoblots revealing the inhibitory effect of *RIPK2/NOD2 KO* on the increase of Lewy-like pathology induced by α-Syn PFF. (E) Bar graph quantifying total α-Syn and p-α-Syn protein levels in the Triton X-100-insoluble fraction in (D) (n=6, each group). Data presented as mean ± SEM (n.s - not significant, ***P < 0.001). (F) Representative photomicrographs showing that *RIPK2/NOD2 KO* blocks the loss of dopaminergic neurons induced by α-Syn PFF in coronal mesencephalon sections containing TH- positive neurons in the SN region. (G) Bar graph quantifying unbiased stereological counts of TH-positive and Nissl-positive neurons in the SN region in (F) (n=8, each group). Data presented as mean ± SEM (n.s - not significant, **P < 0.01, ***P < 0.001). (H) Representative photomicrograph of striatal sections stained for TH immunoreactivity, indicating the genetic depletion of RIPK2/NOD2 rescues α-Syn PFF-induced dopaminergic terminal loss. High-power view of TH fiber density in the striatum. Scale bars: 100 μm (upper panels) and 25 μm (lower panels). (I) Bar graph quantifying dopaminergic fiber densities in (H) (n = 8, each group). Data presented as mean ± SEM (n.s - not significant, **P < 0.01, ***P < 0.001). (J-L) Bar graphs quantifying behavior results, showing that genetic depletion of RIPK2/NOD2 protects against α-Syn PFF-induced motor defects. Results of mice on amphetamine-induced rotation (J), the pole (K), and grip strength (L) (n=13, each group). Data presented as mean ± SEM (n.s - not significant, **P < 0.01, ***P < 0.001).

Consistent with prior reports (Yun et al., 2018), α-Syn PFF-injected *WT* mice showed a substantial loss of TH- and Nissl-positive neurons in the SN, which was significantly attenuated in NOD2^-/-^ or RIPK2^-/-^ mice as assessed by unbiased stereology (Figure 7F and 7G). Western blot analysis revealed that TH and dopamine transporter (DAT) protein levels in the ventral midbrain were reduced in α-Syn PFF-injected *WT* mice, but these reductions were significantly prevented in NOD2^-/-^ or RIPK2^-/-^ mice (S7F and S7G). Moreover, α-Syn PFF-injected *WT* mice showed a significant reduction in striatal TH immunoreactivity (Figure 7H and 7I) and striatal protein levels of TH and DAT (Figure S7H and S7I), while these reductions were attenuated in the striatum of α-Syn PFF-injected NOD2^-/-^ or RIPK2^-/-^ mice (Figure 7H and 7I and S7H and S7I).

To assess motor behavioral deficits in α-Syn PFF-injected mice, we utilized well-established PD-like behavioral tests including the amphetamine rotation, pole, and grip strength tasks as previously described (Brooks and Dunnett, 2009). In α-Syn PFF-injected WT mice, there was an increase in ipsilateral amphetamine-induced rotation, which was significantly prevented in NOD2^-/-^ or RIPK2^-/-^ mice (Figure 7J). Additionally, α-Syn PFF-injected WT showed significant motoric deficits on the amphetamine, pole, and grip strength tasks that were attenuated NOD2^-/-^ or RIPK2^-/-^ mice (Figure 7J and 7L). Collectively, our findings demonstrate that the genetic depletion of NOD2/RIPK2 effectively protects against dopaminergic degeneration and behavior defects in the α-Syn PFF PD mouse model.

## Discussion

The major findings of this study highlight the activation of the NOD2/RIPK2 signaling pathway in microglia as a major contributor to pathological α-Syn induced neurodegeneration. Our findings support the potential of targeting NOD2 and/or RIPK2 activity for therapeutic development to prevent neurodegeneration in PD. This is supported by the findings that genetic NOD2 or RIPK2 depletion in both in vitro and in vivo prevents pathologic α-Syn neuronal injury and reduces the accumulation of pathologic α-Syn.

The innate immune system is vital for immediate defense against harmful stimuli and infections. Recent research has revealed germline-encoded pattern recognition receptors (PRRs) like toll-like receptors (TLRs) and nucleotide-binding oligomerization domain-like receptors (NLRs) such as NOD1 and NOD2. These receptors recognize microbial molecules, such as pathogen-associated molecular patterns (PAMPs), and damage-associated molecular patterns (DAMPs), initiating crucial intracellular signaling pathways for effective defense response in innate immune cells (Heneka et al., 2014; Mogensen, 2009; Takeuchi and Akira, 2010). Microglia, comprising one of the major CNS innate immune cells, are a pivotal component of the CNS immune system. Microglia directly sense damage-associated molecular patterns (DAMPs) or pathogen-associated molecular patterns (PAMPs) via innate immune receptors (Kofler and Wiley, 2011). Mounting evidence suggests the widespread activation of innate immune receptor pathways in neurodegenerative diseases (Heneka et al., 2014; Hirsch and Hunot, 2009).

Misfolded proteins and aggregates in the diseased brain have the potential to serve as DAMPs and trigger neuroinflammation in various brain cells. Our findings strongly support the hypothesis that misfolded and pathologic α-Syn in PD acts as a DAMP. Some studies indicate that TLR2 may be a pattern recognition receptor for higher-ordered oligomeric α-Synuclein (Daniele et al., 2015), or α-Syn PFF (Dutta et al., 2021). However, studies in TLR2-knockout microglia indicate that TLR2 is not required for neurotoxic astrocyte formation (Yun et al., 2018), indicating that other pattern recognition receptors are required for the formation of neurotoxic astrocytes. Our data suggest that α-Syn PFF efficiently serves as DAMPs, activating microglia, with NOD2, a pattern recognition receptor, likely serving as a major receptor for pathological α- Syn in microglia in PD. This hypothesis is substantiated by the following results. First, PLA demonstrates that pathologic α-Syn interacts with the NOD2 receptor in microglia of the substantia nigra in post-mortem PD samples and α-Syn PFF treated WT, but not NOD2^-/-^ mouse microglia. Second, co-immunoprecipitation, surface plasmon resonance (SPR), and indirect ELISA assays indicate that α-Syn PFF interacts with NOD2 in microglia. This interaction leads to the oligomerization of NOD2. Third, human pathologic α-Syn derived from PD patients exhibit a high affinity for human NOD2. Fourth, genetic depletion of NOD2 effectively blocks the downstream signaling pathway initiated by α-Syn PFF interaction with NOD2, both in vitro and in vivo. Fifth, NOD2 mRNA, but not NOD1 mRNA is significantly upregulated in α-Syn PFF activated microglia. Sixth, α-Syn PFF does not bind to either NOD1 or NLRP3, and there is no increase in the colocalization coefficient and area of hNOD1-EGFP and DSRED2-hRIPK2 proteins following α-Syn PFF treatment, further supporting NOD2 as the primary receptor for pathologic α-Syn.

Treatment with α-Syn PFF specifically upregulated NOD2 and RIPK2 protein levels only in microglia, with no significant changes in astrocytes and cortical neurons. In addition, NOD2 and RIPK2 proteins were predominantly observed in microglia, while they were barely expressed in astrocytes and neurons. This was confirmed through PLA, IHC, and WB analyses of the SN from human PD patients and α-Syn PFF-injected mice. Notably, PLA showed that pathologic α-Syn interacts with NOD2 in microglia but not in astrocytes and neurons. Phosphorylation of RIPK2 S176 is thought to be a RIPK2 activation maker (Dorsch et al., 2006; Pellegrini et al., 2017), and consistent with the observation that NOD2 and RIPK2 are primarily expressed in microglia, RIPK2 S176 phosphorylation was only observed in microglia of both PD patients and α-Syn PFF-injected mice. There was an association between NOD2, RIPK2, and phosphorylated α-Syn (pS129 α- Syn) accumulation, indicating a potential pathological relationship. Deletion of NOD2 or RIPK2 reduced pathologic α-Syn accumulation induced by pathologic α-Syn in intrastriatal PFF model as well as enhancement of pathologic α-Syn accumulation by neurotoxic reactive astrocytes suggesting that NOD2 and RIPK2 participate in the accumulation of pathologic α-Syn.

Our research strongly supports the involvement of neurotoxic reactive astrocytes in PD. Previous studies have linked these astrocytes to various neurodegenerative disorders, including glaucoma, ALS, HD, AD, and PD (Guttenplan et al., 2020; Joshi et al., 2019; Liddelow et al., 2017; Sterling et al., 2020; Yun et al., 2018). This underscores their significance as a common response to prolonged CNS injury and highlights their potential as a therapeutic target in neurodegenerative diseases. Emerging evidence suggests that activated microglia can convert astrocytes into neurotoxic reactive forms, harming neurons. This conversion is initiated by the release of three key pro-inflammatory factors: TNFα, IL-1α, and complement C1q, from activated microglia. Notably, regions most affected in PD are rich in neurotoxic reactive astrocytes (Liddelow et al., 2017), further emphasizing the therapeutic potential of targeting this phenomenon in neurodegenerative diseases. Our prior research demonstrated that pathological α-Syn activates microglia, leading to TNFα, IL-1α, and C1q release, driving astrocyte conversion into neurotoxic reactive forms and contributing to dopaminergic neurodegeneration in preclinical PD models (Yun et al., 2018). Building on these findings, our current study identifies key upstream mediators, supporting a crucial role of the NOD2/RIPK2 signaling pathway in connecting activated microglia to neurotoxic reactive astrocyte formation in PD-associated neurodegeneration.

Our work, while insightful, has limitations that will be addressed in future research. First, using global knockouts of NOD2 or RIPK2 inhibitor may impact signaling in other cell types besides microglia. Future studies should employ conditional and inducible strategies to determine if NOD2 or RIPK2 signaling in microglia is essential for the observed effects or if it plays a role in other cells in the CNS like astrocytes and neurons. Additionally, we did not identify the factors secreted by neurotoxic reactive astrocytes in PD and future studies are required to uncover the neurotoxic reactive astrocyte secretome. Addressing these limitations will enhance our understanding of the NOD2/RIPK2 signaling pathway’s therapeutic potential in neurodegenerative diseases, particularly PD. Overall, our findings elucidate the complex interplay among microglia, astrocytes, and neurons in PD pathology. Targeting the NOD2/RIPK2 signaling pathway may hold promise as a therapeutic strategy to mitigate neurotoxic astrocyte activation offering a novel approach for treating neurodegenerative diseases, including PD.

## METHOD DETAILS

### Mouse cortical primary neuron culture

Mouse cortical primary neurons were prepared as described previously (Seo et al., 2018b). Briefly, CD1-timed pregnant mice were purchased from Charles River laboratories (RRID:IMSR_CRL:022). When 15 days pregnant, mice were terminally anesthetized using carbon dioxide (CO_2_) in the euthanasia chamber. The brains were obtained from embryonic day 15 (E15) pups. the meninges were carefully removed, followed by the dissection of the cortex under a dissection microscope. The dissected cortices were then transferred to a 15 ml tube and washed three times using dissection buffer consisting of HBSS (10X, #14185052, Gibco), 100 U/ml Penicillin/streptomycin (10,000 U/ml, #15140122, Gibco), 1 mM sodium pyruvate (100 mM, #11360070, Gibco), 20 mM HEPES (1 M, #15630080, Gibco), and 25 mM Glucose (2.5 M, #G-8769, Sigma). The cortices were incubated in a dissection buffer containing 0.03% trypsin-EDTA (0.25%, #25200056, Gibco) and 50 U/ml DNase (10,000 U, #04536282001, Sigma) in a 37°C water bath for 15 minutes. After incubation, the contents were washed and dissociated in a complete neurobasal medium supplemented with Neurobasal^TM^ medium (#21103-049, Gibco), B- 27 (50X, #17504044, Gibco), Penicillin/streptomycin, and Glutamax-1 (100X, #35050-061, Gibco). The homogenous cells were seeded onto poly-L-lysine-coated coverslip in 12-well plates or 6- well plates, and the cultures were maintained with a complete neurobasal medium. Half of the medium was replaced with fresh medium every 3 days to support the growth and development of cortical neurons. At 3 days in vitro (DIV 3), to eliminate glial cells, the cultures were treated with 5 *μ*M cytosine β-D-arabinofuranoside (AraC, #C1768, Sigma). This protocol can be accessed at protocols.io (https://www.protocols.io/view/mouse-primary-cortical-neuron-culture-c8iszuee; DOI: dx.doi.org/10.17504/protocols.io.e6nvwdr99lmk/v1).

### Mouse primary microglia culture

Primary microglial cultures were performed as described previously (Yun et al., 2018). Brains from postnatal day 1 (P1) pups were collected under dissection microscopy in a petri dish. After carefully removing the meninges, 6 cortices from 3 pups were finely minced into small pieces using spring scissors. The minced cortices were then transferred to a 15 ml tube and washed three times in dissection buffer, which was the same as the one used for mouse primary neuron culture (as mentioned above). The fragmented cortices were incubated in dissection buffer containing 0.125% trypsin-EDTA (2.5%, #15090046, Gibco) and 50 U/ml DNase in a 37°C water bath for 15 min to facilitate digestion. Following the incubation, the cortices were washed three times in DMEM/F12 (#11320033, Gibco) complete medium supplemented with 10% heat-inactivated FBS, Penicillin/streptomycin, L-glutamine (#25030081, Gibco), MEM non-essential amino acids (#11140050, Gibco), and sodium pyruvate (#11360070, Gibco). The DMEM/F12 complete medium was used to halt the trypsinization reaction. A single-cell suspension was obtained by trituration and any cell debris was removed by passing the cell suspension using a 40 μm nylon mesh. The resulting fresh single-cell suspension was seeded on a T175 flask/6 cortices and cultured for 12 days. After seven days of seeding, the medium was changed to maintain cell health and growth. To purify microglia, the EasySep Mouse CD11b Positive Selection Kit II (#18970, StemCell) was used according to the manufacturer’s instructions. The CD11b+ microglia were magnetically separated and cultured separately for further experiments.

### Mouse primary astrocyte culture

Primary astrocytes were isolated by immunopanning and cultured as previously described (Foo et al., 2011; Liddelow et al., 2017). In brief, cortices from P1 mice were diced into 1-mm^3^ pieces and treated with an enzyme solution consisting of 1X EBSS, 0.46% D (+)-Glucose, 26 mM NaHCO_3_, 50 mM EDTA, and distilled H_2_O in the presence of papain, L-cysteine, and DNase. The digested tissues were then repeatedly but gently triturated in the dissociation buffer, also known as low ovomucoid inhibition solution, which contained 1X EBSS, 0.46% D(+)-Glucose, 26 mM NaHCO_3_, distilled H_2_O, 0.002% DNases, 0.10% ovomucoid trypsin inhibitor and 0.10% BSA (Foo et al., 2011). The dissociated cells then went down through the high ovomucoid inhibition solution, which had similar components to the low ovomucoid inhibition solution but with higher concentrations of trypsin inhibitor and BSA to completely block papain activity. The cell suspension was filtered with a pre-soaked Nitex cone containing 0.02% BSA and DNase to remove debris. Subsequently, the cell surface antigens were allowed to recover by incubating the cell suspension at 37°C and 10% CO2 for 30-45 minutes. During this step, microglia, endothelial cells, and oligodendrocytes in the cell suspension were respectively retained on a series of antibody-coated plates, while astrocytes were positively selected on the final anti-ITGβ5 (Integrin beta-5) plate. Following positive selection, the astrocytes were dislodged by trypsinization and cultured in a defined, serum-free base medium containing 50% neurobasal, 50% DMEM, 100 U/ml penicillin, 100 µg/ml streptomycin, 1 mM sodium pyruvate, 292 µg/ml L-glutamine, 1X SATO, 5 µg/ml of N-acetyl cysteine and 5 ng/ml HBEGF (#100-47, Peprotech) as previously described (Liddelow et al., 2017).

### α-Syn purification and α-Syn PFF preparation

Recombinant mouse α-syn monomer proteins were purified using an IPTG independent inducible pRK172 vector system as described previously (Yun et al., 2018). To remove endotoxin, a toxin Eraser endotoxin removal kit (#L00338, Genscript) with a high efficiency endotoxin removal resin was employed. α-syn PFF were prepared as previously described (Volpicelli-Daley et al., 2014). In brief, the endotoxin-removed α-syn monomer was diluted in PBS to a final volume of 500 μl and a concentration of 5 mg/ml. The content was then placed in a Thermomixer set at 1,000 rpm and 37°C for 7 days. After this incubation period, the solution in the tube should appear slightly cloudy. The resulting solution was divided into aliquots (20 μl) each and stored at -80 °C. The quality and structure of α-Syn PFF were validated using atomic force microscopy and transmission electron microscopy. For in vitro treatment, α-Syn PFF at a concentration of 5 mg/ml was added to sterile PBS, resulting in a final concentration of 0.1 mg/ml (20 μl in 980 μl of PBS). The α-Syn PFF was then sonicated with 60 pulses at 10% power, totaling 30 seconds of sonication with 0.5 seconds on and 0.5 seconds off intervals, using a Branson Digital sonifier from Danbury, CT, USA. The sonicated α-Syn PFF was adjusted to a final concentration of 5 μg/ml for use in the target cells. For in vivo injection, the sonicated α-Syn PFF at a concentration of 0.1 mg/ml was stereotaxically injected into the striatum of mice. This protocol is further described here:https://www.protocols.io/view/production-of-synuclein-preformed-fibrils-pff-dm6gpbw28lzp/v1; DOI: dx.doi.org/10.17504/protocols.io.dm6gpbw28lzp/v1).

### Animal

Experimental procedures followed the Laboratory Animal Manual of the National Institute of Health Guide to the Care and Use of Animals and were approved by the Johns Hopkins Medical Institute Animal Care and Use Committee. Animals were housed in a 12-hour dark/light cycle with ad libitum access to water and food. Randomized mixed-gender cohorts were used for all experiments, and mice were acclimatized for 3 days before starting any procedures. Significant efforts were made to minimize animal suffering and discomfort. The α-synuclein pre-formed fibril (α-Syn PFF) induced animal model was established using C57BL/6J mice obtained from the Jackson Laboratories (RRID:IMSR_JAX:000664, Maine, USA). For the present study, these mice were mated with C57BL/6J mice. Six months post-injection, immunohistochemistry and biochemical studies were conducted.

### Human brain tissues

The human post-mortem brain tissues used in this research were obtained with written informed consent from patients, approved by the Johns Hopkins Institutional Review Boards (Approval No. NA00032761). The tissues were sourced from the Morris K. Udall Parkinson’s Disease Research Center of Excellence at Johns Hopkins Medical Institutions (JHMI) and the Alzheimer’s Disease Research Center (http://www.alzresearch.org), in accordance with local Institutional Review Board and HIPAA (Health Insurance Portability and Accountability Act) regulations.

### RNA sequencing analysis for transcriptome strand-specific

The total RNA from primary microglia treated with α-syn PFF 5ug/ml undergoes rRNA removal using the Ribo-Zero kit (#20040526; Illumina). Subsequently, the remaining RNA is purified, fragmented, and primed for cDNA synthesis. The cleaved RNA fragments are converted into first-strand cDNA using reverse transcriptase and random hexamers, followed by second-strand cDNA synthesis with DNA Polymerase I, RNase H, and dUTP. These cDNA fragments undergo end repair, have a single ’A’ base added, and are ligated with adapters. The resulting products are purified and enriched with PCR to generate the final cDNA library. Illumina employs a unique "bridged" amplification reaction on the flow cell’s surface for clustering and sequencing. The HiSeq 2000 automates cycles of extension and imaging with a flow cell containing millions of distinct clusters. Solexa’s Sequencing-by-Synthesis method utilizes four proprietary nucleotides with reversible fluorophore and termination properties. Each sequencing cycle incorporates all four nucleotides simultaneously, resulting in higher accuracy compared to methods with a single nucleotide in the reaction mix at a time. This cycle is iterated one base at a time, generating a series of images, each representing a single base extension at a specific cluster. To assess sequencing quality, FastQC is utilized for quality control checks on raw sequence data from high-throughput sequencing pipelines. It provides a modular set of analyses to quickly identify any potential issues before further analysis. For data analysis, TopHat, a fast splice junction mapper, aligns RNA-Seq reads to mammalian-sized genomes using Bowtie, an ultra-high-throughput short read aligner. It then identifies splice junctions between exons based on the mapping results. Cufflinks are used for assembling transcripts, estimating their abundances, and testing for differential expression and regulation in RNA-Seq samples. It accepts aligned RNA-Seq reads and assembles the alignments into a parsimonious set of transcripts. Cufflinks estimates the relative abundances of these transcripts, accounting for biases in library preparation protocols. Further information for FastQC can be found here: http://www.bioinformatics.babraham.ac.uk/projects/fastqc (version: v0.10.0). More information on TopHat can be found here: http://tophat.cbcb.umd.edu/index.html (version: TopHat v2.0.11, Bowtie2 version 2.1.0). For Cufflinks, additional details are available here: http://cufflinks.cbcb.umd.edu/index.html (version: cufflinks v2.1.1).

### Bioinformatics analysis of RNA-seq

Differential expression analysis for raw RNA-seq counts was performed using R 4.3.2 (RRID:SCR_000432, www.r-project.org). Raw counts were imported into the R Studio environment using the EdgeR package (Version 4.0.2) for downstream analysis. To account for variations in sequencing depth and composition biases between samples, TMM normalization was performed using the EdgeR package. Raw counts were transformed into log-Counts Per Million (log-CPM) values using the boom function from the Limma package (Version 3.58.1). A standard linear model was fitted to the log-CPM values using the lmFit function from the limma package. Empirical Bayes moderation of the model fit was achieved through the eBayes function. Differential expression analysis was conducted using the results from the limma model fit. The topTable function was employed to extract differentially expressed genes based on the specified contrast of interest. Gene Ontology (GO) analysis and Kyoto Encyclopedia of Genes and Genomes (KEGG) pathway analysis were employed to identify gene functions and molecular pathways of differentially expressed genes (DEGs) in α-syn PFF-activated microglia compared to non-activated microglia. The DEGs were submitted to the online software Database for Annotation, Visualization, and Integrated Discovery (DAVID) for GO pathway analyses and KEGG pathway analyses. Enrichment analyses for GO terms and KEGG pathways were conducted using the (DAVID) online database (Bioinformatics Resources 2021, RRID:SCR_001881DAVID).

### Gene ontology and pathway enrichment analyses of DEGs

Gene Ontology (GO) analysis was employed to identify gene functions and molecular pathways of differentially expressed genes (DEGs) in α-syn PFF-activated microglia compared to non-activated microglia. The DEGs were submitted to the online software DAVID for GO pathway analyses. Enrichment analyses for GO terms were conducted using the Database for Annotation, Visualization, and Integrated Discovery (DAVID) online database (DAVID Bioinformatics Resources 6.8, RRID:SCR_001881DAVID).

### In situ proximity ligation assay (PLA)

The duolink proximity ligation assay was used to visualize the interaction between pathologic α- syn aggregates and endogenous NOD2 protein and performed as described previously (Seo et al., 2021; Soderberg et al., 2006). Primary microglia were purified and seeded onto poly-L-lysine-coated coverslips in a 12-well plate (1×10^5^ cells/well). After 1 day, the cells were treated with α- Syn PFF for 1 hour. For the assay, the cells were washed three times with PBS and fixed using 4% paraformaldehyde with 4% sucrose. Permeabilization was performed with 0.2% Triton X-100 in PBS, and then the in situ proximity ligation assay was carried out following the manufacturer’s instructions (#DUO92101, Sigma-Aldrich). The cells were blocked with PLA blocking buffer and incubated with anti-NOD2 antibody (#MA1-16611, ThermoFisher, 1:500: RRID: AB_568643), anti-α-syn aggregate antibody (#ab209538, Abcam, 1:500; RRID:AB_2714215), and anti-Iba1 antibody (#PA5-18039, ThermoFisher, 1:500 for visualizing microglial cells, RRID: AB_10982846) at 4°C for 1 hour in a humidity chamber. Minus- or Plus-probe conjugated secondary antibodies were then applied and incubated at 37°C for 1 hour in a preheated humidity chamber. Subsequently, the ligation mix was added to each coverslip and incubated at 37°C for 30 minutes in a preheated humidity chamber. The signals were amplified by adding an amplification-polymerase containing reaction solution and incubating at 37°C for 100 minutes in a preheated humidity chamber. The coverslips were mounted with a DAPI-containing mounting medium, and the images were captured by LSM880 confocal microscopy. The number of positive signals (red signals) was quantified using ImageJ software (RRID:SCR_003070).

### In vitro protein binding assay

In the NeutrAvidin full-down assay (Figure S2A), plasmids expressing hNOD1, hNOD2, hNLRP3, and hRIPK2 with EGFP were transfected into HEK 293T cells. After 48 hours, cells were treated with biotin-conjugated α-Syn PFF or α-syn monomer and lysed using immunoprecipitation lysis buffer (#87787, Thermo). Cell lysates were then incubated with neutravidin (#29201, Thermo) to explore α-Syn-biotin PFF bound to the beads. The proteins precipitated with α-Syn-biotin PFF or monomer through immunoblotting with anti-GFP.

In the reciprocal full-down assay (Figure S2D), HEK293T cells expressing His-Myc-tagged FL hNOD2 or various deletion mutants (ΔCARD, ΔNBD, and ΔLRR) were treated with α-Syn-biotin PFF or α-Syn-biotin monomer and lysed. Cell lysates were incubated with neutravidin or nickel-NTA agarose beads (#A50585, Thermo) to explore α-Syn-biotin PFF and α-Syn-biotin monomer bound to the beads. Immunoblotting with anti-myc was performed to analyze the proteins precipitated with α-Syn-biotin PFF or monomer.

In the reciprocal pull-down assays (Figure S4A), a plasmid expressing His-Myc-FL hNOD2 was transfected into BV2 cells. After 48 hours, α-Syn PFF was treated, and cells were collected at indicated time points. Anti-Myc antibody or NeutrAvidin was used for immunoprecipitation of Myc-tagged hNOD2 or to pull down α-Syn-biotin PFF, respectively.

### Immunocytochemistry and image analysis

For the immunostaining assay in Figure S4F, BV2 cells were plated and grown on 10-mm coverslips. Plasmids expressing hNOD1-EGFP/DSRED2-hRIPK2 or hNOD2-EGFP/DSRED2- hRIPK2 were co-transfected into BV2 cells. After 48 hours, the cells were treated with α-Syn PFF and then washed three times with PBS. They were fixed using 4% paraformaldehyde (#15710, Electron Microscopy Science) at the indicated time points. The paraformaldehyde-fixed cells were permeabilized with 0.2% Triton X-100 in PBS and washed three times with PBS. The cells were then mounted using gold antifade reagent (#P36930, ThermoFisher), and fluorescence was visualized using a confocal microscopy system with a ZEIZZ LSM880 microscope equipped with a 60x oil-immersion objective. The images were processed with the LSM Image Software (Zeiss LSM software (RRID:SCR_014340). The Manders colocalization coefficient was analyzed using ImageJ, specifically JACoP. Manders’ M1 and M2 coefficients were calculated to determine the degree of colocalization between a pair of proteins stained and imaged in two channels (red and green) (Bolte and Cordelieres, 2006). Manders M1 determines the degree of green channel colocalizing with red, and M2 determines the reverse. For the immunocytochemistry assay, cells used in this study were fixed with freshly prepared paraformaldehyde (4%) for 15 minutes and washed three times with artificial cerebrospinal fluid (ACSF; 0.15 M NaCl, 10 mM HEPES, 3 mM KCl, 0.2 mM CaCl2 dihydrate, 10 mM glucose). Fixed cells were permeabilized with 0.2% Triton X-100 in PBS before adding a blocking buffer (containing 3% BSA and 0.1% Triton X-100 in PBS) for an hour at room temperature. Cells were then incubated with the primary antibody in a blocking buffer (1:500) for an hour at 4°C. Next, a secondary Alexa Fluor® antibody (1:1000) was added to each well for 2 hours, followed by DAPI staining for 5 minutes (#H-1200-NB, Vectashield®). Cells were washed with PBS (containing 0.1% Triton X-100) three times between incubations. The images were captured using LSM880 confocal microscopy. Immunostaining protocol can be found here: https://www.protocols.io/view/immunostaining-c8j5zuq6; DOI: dx.doi.org/10.17504/protocols.io.81wgbxb73lpk/v1).

### Blue native-polyacrylamide gel electrophoresis (BN-PAGE)

For the immunoblot assay in Figure S4J, primary microglia grown on 10-mm coverslips were transduced with AAV expressing hRIPK2 with P2A (cleavage site) and EGFP or lentivirus carrying HA (human influenza hemagglutinin)-tagged hNOD2. After 5 days, α-Syn PFF (5 ug/ml) was treated, and cells were collected at indicated time points. BN-PAGE was utilized to determine the oligomeric state between NOD2 and RIPK2. Primary microglia expressing hNOD2/hRIPK2 were lysed using a non-ionic (NP-40) lysis buffer to extract cytoplasmic proteins at specified time points. Native PAGE was then conducted as previously described (Abdel-Nour et al., 2019).

### Protein expression and purification Fc-fused hNOD2 FL, ΔCARD, ΔNBD, and ΔLRR

Fc-fused human NOD2 FL, ΔCARD, ΔNBD, and ΔLRR were purified from HEK293T cells (RRID: CVCL_0063) using a transient transfection protocol as described previously (Seo et al., 2018a). Briefly, cells were cultured in T175 cm2 plates to 85–90% confluency and transfected with the target vectors (10 µg/plate) using TransIT-X2® dynamic delivery transfection reagent (#MIR6006, Mirus). After 48 hours post-transfection, the cells were harvested via centrifugation at 6,000 rpm, and the cell pellet was re-suspended and lysed by sonication in a NOD2 protein purification buffer containing 25 mM Tris pH 8.0, 500 mM NaCl, 10% glycerol, and 1 mM DTT as described previously (Maekawa et al., 2016). Additionally, 0.1mM ADP was included in all buffers during the purification process. The lysate was then subjected to ultracentrifugation at 13,000 rpm and 4°C for 2 hours, and the supernatant was filtered through a 0.2-µm syringe filter (Millipore). To reduce non-specific binding and background, the filtered supernatant was pre-incubated with Sepharose beads (6B100, Sigma) and subsequently incubated with Protein A Sepharose beads (#17-0780- 01, GE Healthcare). After washing with a NOD2 protein purification buffer containing 0.1% Triton X-100, 0.5 mM EDTA, and 0.5 mM EGTA, the target proteins were eluted using an elution buffer containing 0.2 M glycine (pH 2.5). Proteinase inhibitors and ADP were added to all purification steps. Purified proteins were then subjected to a buffer change using a storage buffer containing 20 mM Tris pH 8.0, 150 mM NaCl, and 1% glycerol, and stored at -20 °C.

### Indirect ELISA

Fc-fused NOD2 FL and deletion mutants were generated in HEK293T cells and purified using protein A-beads. α-syn PFF, α-syn monomer, or BSA (50 μg/ml) were coated in a 96-well maxisorp plate (#44-2404-21, Thermo) and then incubated with various concentrations of Fc-fused NOD2 FL and deletion mutants. To detect NOD2 proteins binding to α-Syn PFF or monomer, the plate was washed 5 times with PBS and incubated for 1 hour with goat anti-human IgG (Fc specific) –peroxidase (#A0170, Sigma). Binding signals were developed and stopped by the addition of TMB (3,3’,5,5’-tetramethylbenzidine) solution (#T0440, Sigma) and sulfuric acid, and the colorimetric reaction was evaluated by measuring the absorbance at 450 nm. For a competition ELISA, NOD2 was preincubated with α-Syn PFF or α-Syn monomer, a competitor, for 30 minutes before its addition to the α-Syn PFF- or α-Syn monomer-coated plate. All binding was quantified by measuring the activity of Fc antibody-coupled peroxidase.

### Cleavage of Fc tag from Fc-fused hNOD2 FL

To determine the precise binding kinetics between NOD2 and α-syn PFF, a 50 µg sample of Fc-fused hNOD2 FL protein containing an HRV 3C cleavage site (human NOD2-HRV 3C-Fc) was incubated with 1 µg of HRV 3C protease (#88946, Thermo) at a 50:1 (w/w) ratio in 500 µl of 20 mM Tris pH 8.0, 150 mM NaCl, and 1% glycerol at 4 °C. The HRV 3C protease was fused with a GST-tag (glutathione-S-transferase) to facilitate its removal from the samples after cleavage. After a 16-hour incubation period, the reaction solution was sequentially precipitated using protein A sepharose beads and Glutathione Sepharose beads (#17-0756-01, GE Healthcare). The resulting supernatant was stored at -20 °C until it was analyzed for the surface plasmon resonance (SPR) experiment.

### Surface Plasmon Resonance (SPR)

To investigate the real-time interaction between α-Syn aggregates and NOD2, we conducted binding measurements and kinetics analysis using surface plasmon resonance (SPR) on an open SPR instrument (Nicoya). Specifically, α-Syn PFF or α-Syn monomer, along with human patient- or healthy control-derived PMCA products, were immobilized at a high coupling density on high sensitivity carboxy sensor chips in the active flow cell (channel 2). Subsequently, we passed increasing concentrations (ranging from 125 nM to 3 μM) of NOD2 FL and domain mutants over both the active flow cell (channel 2) and an uncoated reference flow cell (channel 1). To determine the interaction affinity, we utilized both bind kinetics and steady-state affinity measurements. The sensorgrams obtained were then analyzed using the TraceDrawer software, which allowed us to extract real-time binding kinetics and affinity data from the experiments.

### α-Synuclein Protein misfolding cyclic amplification (α-Syn PMCA)

The α-Syn-PMCA assay was performed as described previously (Shahnawaz et al., 2020) with some modifications using the PMCA equipment from Qsonica (Qsonica, Newtown, CT, USA). In brief, full-length human α-Syn (residues 1–140) was expressed and purified from Escherichia coli strainBL21(DE3). The monomeric α-Syn was then centrifuged at 100,000 g for 30 minutes at 4 °C to eliminate any preformed aggregates. The resulting seed-free α-syn was diluted in PMCA buffer (1% Triton X-100 in PBS) at a concentration of 0.1 mg/mL and placed in PCR tubes with a final volume of 100 μl. For each PCR tube, 10 μl of cerebrospinal fluid (CSF) from both controls and PD patients (Table S1) was added as seeds, and this setup was performed in triplicate. Silicon beads with a diameter of 1.0 mm were introduced into the PCR tubes before adding the mixture. The samples were subjected to the PMCA reaction, which involved 20 seconds of sonication (at 10% amplitude) followed by 29 minutes and 40 seconds of incubation at 37 °C for 7 days.

### Thioflavin T (ThT) fluorescence assay

The degree of α-syn fibrillation was continuously monitored by Thioflavin T (ThT) fluorescence, which exhibits an increase in fluorescence when it binds to amyloid-like fibrils. Each day, a 6 µl sample was withdrawn from each PCR tube and diluted in 150 µl of ThT solution (final ThT concentration 20 µM). The samples were thoroughly mixed at room temperature, and then 50 µl of each diluted sample was transferred to a black clear-bottom 384-well plate. After incubating the plate at 37 °C for 5 minutes, the fluorescence intensity was measured at an excitation wavelength of 450 nm and emission wavelength of 485 nm using a Varioskan™ LUX multimode microplate reader. See protocol here: https://www.protocols.io/view/thioflavin-t-fluorescence-assay-c8kbzusn; DOI: dx.doi.org/10.17504/protocols.io.n2bvj346xlk5/v1)

### Transmission electron microscopy (TEM)

The PMCA samples were diluted 50-fold in double distilled water and carefully adsorbed onto 400 mesh carbon-coated copper grids obtained from Ted Pella Inc., USA. Any excess buffer was removed using lint-free filter paper. Next, the samples underwent negative staining by applying 2% uranyl acetate for 1 minute, followed by gentle drying with lint-free filter paper. The prepared grids were then examined and imaged using a Hitachi H7600 electron microscope equipped with AMT Cameras (M088 Smith), employing an acceleration voltage of 80 kV.

### Immunoblotting for PMCA products

PMCA products from control and PD patients were analyzed by Western blot. The samples (2 μg) were loaded onto a 15% SDS-PAGE gel and transferred to PVDF membranes (0.2 μm, Bio-Rad). After blocking with 5% BSA in TBST at room temperature for 1 hour, the membrane was incubated with an α-Syn-specific primary antibody (BD Transduction, USA) at a dilution of 1:1000 overnight at 4°C. Subsequently, the membrane was treated with secondary antibodies (Millipore Sigma) at a dilution of 1:5000 for 2 hours at room temperature. Image detection was performed using an Amersham Imager 600 (GE Healthcare Life Sciences, USA).

### Nuclear and cytoplasmic extraction

Cultured primary microglia were fractionated into nuclear and cytoplasmic fractions using NE-PERTM Nuclear and cytoplasmic extraction reagents (#78833, Thermo). The fractions were confirmed by detecting heat shock protein 90 (Hsp90) in the cytoplasmic fraction and lamin A/C in the nuclear fraction.

### TUBE pull-down assay

TUBE pulldown was performed following the manufacturer’s instructions (#UM411, Lifesensors). At the specified time points, WT, RIPK2^-/-^, and NOD2^-/-^ primary microglia treated with M18-MDP (1 μg/ml) or α-Syn PFF (5 μg/ml) were lysed in ubiquitination lysis buffer (#PI89900, ThermoFisher), supplemented with protease inhibitor, 50 μM PR-619 (#SI9619, Lifesensors), and 5 mM 1,10-phenanthroline (o-PA) (100X, #SI9649, LifeSensors). The total protein content of the lysates was determined by BCA protein assay (#23227, ThermoFisher), and 20 µl of Anti-Ub TUBE agarose resin was added. The reactions were rotated for 2 hours at 4°C, washed three times with TBST, and the beads were resuspended in 100 µL of elution wash buffer. After incubating for 5 minutes at room temperature, the beads were resuspended in 50 µL of elution buffer. The supernatant was neutralized with 10x Neutralization buffer and analyzed by SDS-PAGE and immunoblotting for the target proteins.

### Stereotaxic α-Syn PFF injection into mouse striatum

For stereotaxic injection of α-Syn PFF into the mouse striatum, 3-month-old male and female mice were anesthetized with 2% avertin (Sigma) and positioned in a stereotaxic apparatus (David Kopf instruments). A glass pipette (diameter, 15–20 µm) was stereotaxically inserted into the striatum unilaterally (right hemisphere) at coordinates mediolateral 2.0 mm from bregma, anteroposterior -0.2 mm, and dorsoventral 2.6 mm. A total of 1 µl of α-Syn PFF solution (0.1 mg/ml) or an equivalent volume of PBS was injected using a picospritzer, and the inserted glass pipette was left in place for at least 10 minutes post-infusion. Full protocol can be accessed here: https://www.protocols.io/view/stereotaxic-syn-pff-injection-into-mouse-striatum-c8iwzufe; DOI: dx.doi.org/10.17504/protocols.io.36wgq3853lk5/v1).

### Immunohistochemistry and image analysis

*In vivo* immunohistochemistry was performed as described previously (Kwon et al., 2021) and was perfused with ice-cold PBS, followed by 4% paraformaldehyde. Brains were dissected and post-fixed overnight in 4% paraformaldehyde at 4°C and then cryoprotected in 30% sucrose solution. Serial brain sections, 30 μm thick, were cut from the fixed brains. The sections were blocked in 5% goat or horse serum (#005-000-121, #008-000-121, Jackson ImmunoResearch) with 0.2% Triton X-100 and incubated with anti-pS129-α-syn antibodies (#ab184674, Abcam, 1:500, RRID:AB_2819037), anti-TH (#NB300-109, Novus Biologicals, 1:500, RRID:AB_10077691), anti-Iba-1 (#019-19741, Wako, RRID:AB_839504), or anti-GFAP (#Z03340-2, Agilent, 1:500). Subsequently, biotin-conjugated anti-rabbit (1:250) or mouse (1:250) antibodies were used, and the sections were washed three times. The prepared sections were then incubated with ABC reagents and developed using SigmaFast DAB Peroxidase Substrate (D4293, Sigma-Aldrich). Additionally, the sections were counterstained with Nissl (0.09% thionine). Image analysis was performed using a computer-assisted system, including an Axiophot photomicroscope (Carl Zeiss) with a motorized stage (Ludl Electronics, NY, USA), a Hitachi HV C20 camera, and Stereo Investigator software (MicroBright-Field, VT, USA). TH- and Nissl-positive dopaminergic neurons in the SN region were counted blindly by an investigator. The total number of TH-stained neurons and Nissl counts were evaluated as described previously (Kwon et al., 2021). Additionally, the fiber density in the striatum was analyzed using optical density measurement. For the analysis of pS129-α-Syn, Nissl-positive cells, microglia, and astrocytes in the SN region, ImageJ software and field counting methods were utilized. In the case of paraffin-embedded human brain tissues, slides were deparaffinized in xylene, rehydrated in ethanol, and washed in distilled water before the application of the primary antibody. Antigen retrieval was performed at 90°C for 20 minutes using sodium citrate buffer, pH 6 (10X, #ab64214, Abcam). Subsequently, slides were washed in 1x TBST, and endogenous peroxidases and alkaline phosphatases were blocked using BLOXALL® Endogenous Blocking Solution (#SP- 6000-100, Vector Laboratories) for 10 minutes, followed by another rinse in 1x TBST. Full protocol can be seen here: https://www.protocols.io/view/immunohistochemistry-and-image-analysis-c8izzuf6; DOI: dx.doi.org/10.17504/protocols.io.ewov1q682gr2/v1).

### Immunofluorescence analysis and image analysis

Brain tissue sections, 30 μm thick, were prepared using a Leica cryostat. The following antibodies were applied: anti-pS129-α-Syn antibodies (#ab184674, Abcam, 1:500), anti-TH (#NB300-109, Novus Biologicals, 1:500), anti-Iba-1 (#019-19741, Wako), or anti-GFAP (#Z03340-2, Agilent, 1:500). Visualizations of the primary antibodies were achieved using suitable secondary antibodies conjugated with Alexa fluorophores (Alexa Fluor 488, 594, and 647, Invitrogen).

### Preparation of PFF-MCM and PFF-ACM

The activated conditioned medium was prepared as described previously (Yun et al., 2018). The activated conditioned medium was prepared following previous methods (Yun et al., 2018). Conditioned medium from primary microglia incubated with α-Syn PFF (PFF-MCM) with either PBS or CMPD0673 treatment was collected and transferred to primary astrocytes for 24 hours. The conditioned medium from activated astrocytes by PFF-MCM (PFF-ACM) was collected and concentrated using an Amicon Ultra-15 centrifugal filter unit (10 kDa cutoff, #UPC9010, Millipore) to approximately 50-fold concentration. Total protein concentration was measured using the Pierce BCA protein assay kit, and 15 μg/ml of total protein was added to mouse primary neurons for the neuronal cell death assays.

### Purification of astrocytes and microglia from adult mice

Astrocytes and microglia from the ventral midbrain of adult mice were isolated as described previously (Batiuk et al., 2017). Each experiment involved 3 mice (6 months after α-syn PFF-injection). Dissected ventral midbrains were dissociated using the Neural Tissue Dissociation Kit (P, papain) (#130-092-628, Miltenyi Biotec) and Neural Tissue Dissociation Kit (T, trypsin) (#130- 093-231, Miltenyi Biotec) following the manufacturer’s instructions. After the enzymatic reaction, cells were mechanically triturated with 10 ml serological pipettes (3 rounds of 10 strokes each). The dissociated cells were then passed through a 20 μm Nitex filter to remove remaining tissue clumps. Myelin was further removed using equilibrium density centrifugation with 90% Percoll PLUS (#GE17-5445-01, Cytiva) in 1× HBSS with calcium and magnesium (#55037C, Sigma). The final Percoll concentration was adjusted to 24%. DNase I (Worthington) was added (1250 units per 10 ml of suspension), and the suspension was gently mixed before being spun down at 300g for 10 minutes at room temperature. The cell-containing pellet was resuspended in 0.5% BSA in PBS without calcium and magnesium. For the isolation of adult microglia and astrocytes, an additional myelin removal step was performed using Myelin Removal Beads II (#130-096-731, Miltenyi Biotec) following the manufacturer’s instructions, and LS magnetic columns (#130-042- 401, Miltenyi Biotec) were used. The flow-through fraction was collected for subsequent astrocyte isolation. For positive selection of astrocytes and microglia, the ACSA-2 kit (#130-097-678, Miltenyi Biotec) and CD11b (#130-093-634, Miltenyi Biotec) were used, respectively, following the manufacturer’s instructions. Two runs of enrichment were performed on consecutive MS columns (#130-042-201, Miltenyi Biotec).

### Microelectrode array (MEA) recordings and data analyses

Cortical cultured neurons were plated on a single-well MEA plate with 64 channels and Pt recording electrodes (Axion biosystem, U.S.). The recording electrodes had a diameter of 30 μm and a spacing of 200 μm center to center. The plate was placed on the Muse MEA system chamber (Axion biosystem, U.S.), and data were acquired and analyzed using the Axion integrated studio software (AxiS) at a sampling frequency of 25 kHz. For spike sorting, raw data were filtered offline at 200 Hz and 3000 Hz using a Butterworth high-pass and low-pass filter. Spike cutouts were then detected offline based on an adaptive threshold crossing to 6 x standard deviations through the spike detector (AxiS) and transferred to Neural Metric Tool software (Axion biosystem). Spikes were plotted in 2-D. The recording was performed at DIV 7, DIV 10, and DIV 14 after cell plating, continuously under 37°C using the control module. Mean firing rates were calculated with all units over the time scale.

## ACKNOWLEDGEMENT

All relevant ethical regulations were followed. This work was supported by grants from the NIH/NINDS NS107404 (H.S.K.), and a sponsored research agreement with Neuraly, the JPB Foundation (T.M.D.), RF1 NS125592 (X.M.), NS38377 (T.M.D., H.S.K., L.S.R.), and PF-PRF-1046133 (K.G). T.M.D. is the Leonard and Madlyn Abramson Professor in Neurodegenerative Diseases. We thank Dr. Santanu Bose at Department of Microbiology and Immunology for NOD2 depletion mutants, The University of Texas Health Science Center at San Antonio for the generous gifts of cDNA plasmids for NOD2. For the purpose of open access, the author has applied a CC-BY public copyright license to the Author Accepted Manuscript (AAM) version arising from this submission.

## Author Contributions

B.A.S. and S.K. designed a majority of the experiments, performed the experiments, analyzed data, and wrote the manuscript. B.A.S., S.K., and D.K. performed experiments and data interpretation. H.S.K., B.A.S., S.K., and S.M. performed sample preparation and helped with experiments. B.A.S. and S.K. provided and managed mice and PFF and helped with data interpretation. S.L. helped with data interpretation. K.G., X.M, and L.S.R. contributed to PMCA work. H.S.K., B.A.S., and S.K., T.M.D, and V.L.D. performed manuscript writing, review, and editing. J.C.T., and J.R. provided human postmortem brain samples. N.B. provided illustrate images. H.S.K. supervised the project, formulated the hypothesis, designed experiments, analyzed data, and wrote the manuscript.

## Competing Financial Interests Statement

H.S.K., T.M.D, and V.L.D. are co-founders of Neuraly and hold equity in D&D Pharmatech, a holding company of Neuraly. This arrangement has been reviewed and approved by the Johns Hopkins University in accordance with its conflict of interest policies. H.S.K. has received payments as a consultant and funding from Neuraly. S.L. is an employee of Neuraly.

---

**Table S1.**
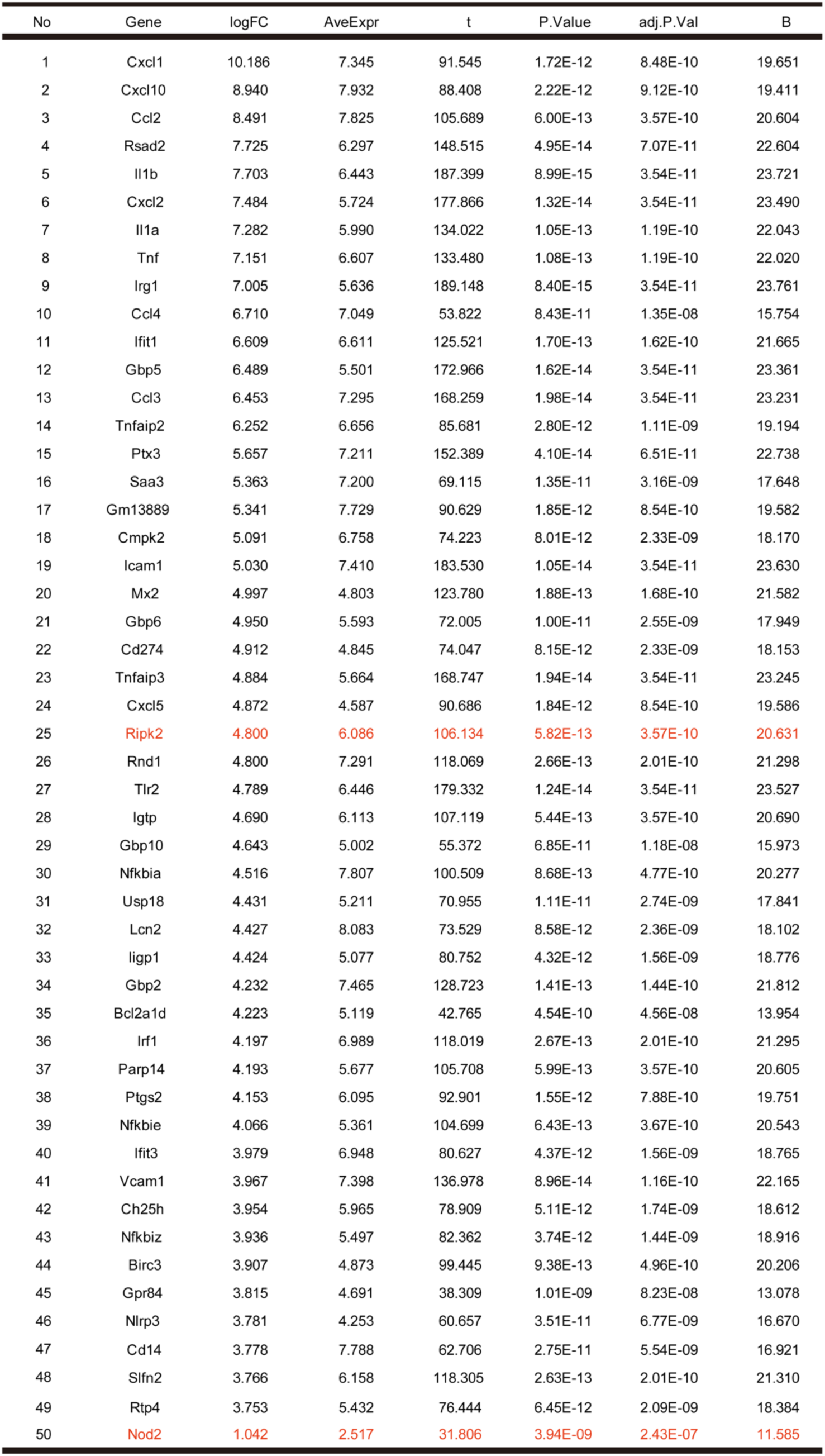
Differentially Expressed Genes by RNA-Seq data analysis.

**Table S2.** Human post-mortem brain tissues and cerebrospinal fluid (CSF) used in Figure 2A and Figure S2Q, respectively.

| Age | Sex | Race | Diagnosis | Region | Immunoblotting | Immunohistochemistry (IHC) | α-syn PMCA |
| --- | --- | --- | --- | --- | --- | --- | --- |
| 73 | F | W | Control | Substantia nigra | 0 |  |  |
| 79 | M | W | Control | Substantia nigra | 0 |  |  |
| 79 | M | W | Control | Substantia nigra | 0 | 0 |  |
| 89 | M | W | Control | Substantia nigra | 0 | 0 |  |
| 68 | F | W | Control | Substantia nigra | 0 |  |  |
| 87 | F | W | Control | Substantia nigra | 0 | 0 |  |
| 65 | F | W | Control | CSF |  |  | 0 |
| 69 | F | W | Control | CSF |  |  | 0 |
| 82 | F | W | Control | CSF |  |  | 0 |
| 71 | M | W | Control | CSF |  |  | 0 |
| 67 | F | W | Control | CSF |  |  | 0 |
| 76 | M | W | PD with Dementia | Substantia nigra | 0 |  |  |
| 83 | F | W | PD with Dementia | Substantia nigra | 0 | 0 |  |
| 83 | M | W | PD with Dementia | Substantia nigra | 0 | 0 |  |
| 84 | F | W | PD with Dementia | Substantia nigra | 0 | 0 |  |
| 80 | F | W | PD with Dementia | Substantia nigra | 0 |  |  |
| 89 | M | W | PD with Dementia | Substantia nigra | 0 |  |  |
| 78 | M | W | PD with Dementia | CSF |  |  | 0 |
| 68 | M | W | PD with Dementia | CSF |  |  | 0 |
| 79 | M | W | PD with Dementia | CSF |  |  | 0 |
| 66 | M | W | PD with Dementia | CSF |  |  | 0 |
| 74 | M | W | PD with Dementia | CSF |  |  | 0 |

## Notes

### Competing Interest Statement

The authors have declared no competing interest.

### Summary of Updates

The grant information is revised on ACKNOWLEDGEMENT.

## Reference

Abdel-Nour, M., Su, H., Duncan, C., Li, S., Raju, D., Shamoun, F., Valton, M., Ginevra, C., Jarraud, S., Guyard, C., et al. (2019). Polymorphisms of a Collagen-Like Adhesin Contributes to Legionella pneumophila Adhesion, Biofilm Formation Capacity and Clinical Prevalence. Front Microbiol 10, 604.

Abounit, S., Bousset, L., Loria, F., Zhu, S., de Chaumont, F., Pieri, L., Olivo-Marin, J.C., Melki, R., and Zurzolo, C. (2016). Tunneling nanotubes spread fibrillar alpha-synuclein by intercellular trafficking of lysosomes. EMBO J 35, 2120–2138.

Anderson, M.A., Burda, J.E., Ren, Y., Ao, Y., O’Shea, T.M., Kawaguchi, R., Coppola, G., Khakh, B.S., Deming, T.J., and Sofroniew, M.V. (2016). Astrocyte scar formation aids central nervous system axon regeneration. Nature 532, 195–200.

Ashburner, M., Ball, C.A., Blake, J.A., Botstein, D., Butler, H., Cherry, J.M., Davis, A.P., Dolinski, K., Dwight, S.S., Eppig, J.T., et al. (2000). Gene ontology: tool for the unification of biology. The Gene Ontology Consortium. Nat Genet 25, 25–29.

Batiuk, M.Y., de Vin, F., Duque, S.I., Li, C., Saito, T., Saido, T., Fiers, M., Belgard, T.G., and Holt, M.G. (2017). An immunoaffinity-based method for isolating ultrapure adult astrocytes based on ATP1B2 targeting by the ACSA-2 antibody. J Biol Chem 292, 8874–8891.

Bolte, S., and Cordelieres, F.P. (2006). A guided tour into subcellular colocalization analysis in light microscopy. J Microsc 224, 213–232.

Borst, K., Dumas, A.A., and Prinz, M. (2021). Microglia: Immune and non-immune functions. Immunity 54, 2194–2208.

Brooks, S.P., and Dunnett, S.B. (2009). Tests to assess motor phenotype in mice: a user’s guide. Nat Rev Neurosci 10, 519–529.

Choi, Y.R., Park, S.J., and Park, S.M. (2021). Molecular events underlying the cell-to-cell transmission of alpha-synuclein. FEBS J 288, 6593–6602.

Chou, S.C., Aggarwal, A., Dawson, V.L., Dawson, T.M., and Kam, T.I. (2021a). Recent advances in preventing neurodegenerative diseases. Fac Rev 10, 81.

Chou, T.W., Chang, N.P., Krishnagiri, M., Patel, A.P., Lindman, M., Angel, J.P., Kung, P.L., Atkins, C., and Daniels, B.P. (2021b). Fibrillar alpha-synuclein induces neurotoxic astrocyte activation via RIP kinase signaling and NF-kappaB. Cell Death Dis 12, 756.

Clausson, C.M., Allalou, A., Weibrecht, I., Mahmoudi, S., Farnebo, M., Landegren, U., Wahlby, C., and Soderberg, O. (2011). Increasing the dynamic range of in situ PLA. Nat Methods 8, 892–893.

Damgaard, R.B., Nachbur, U., Yabal, M., Wong, W.W., Fiil, B.K., Kastirr, M., Rieser, E., Rickard, J.A., Bankovacki, A., Peschel, C., et al. (2012). The ubiquitin ligase XIAP recruits LUBAC for NOD2 signaling in inflammation and innate immunity. Mol Cell 46, 746–758.

Daniele, S.G., Beraud, D., Davenport, C., Cheng, K., Yin, H., and Maguire-Zeiss, K.A. (2015). Activation of MyD88-dependent TLR1/2 signaling by misfolded alpha-synuclein, a protein linked to neurodegenerative disorders. Sci Signal 8, ra45.

Dorsch, M., Wang, A., Cheng, H., Lu, C., Bielecki, A., Charron, K., Clauser, K., Ren, H., Polakiewicz, R.D., Parsons, T., et al. (2006). Identification of a regulatory autophosphorylation site in the serine-threonine kinase RIP2. Cell Signal 18, 2223–2229.

Dutta, D., Jana, M., Majumder, M., Mondal, S., Roy, A., and Pahan, K. (2021). Selective targeting of the TLR2/MyD88/NF-kappaB pathway reduces alpha-synuclein spreading in vitro and in vivo. Nat Commun 12, 5382.

Foo, L.C., Allen, N.J., Bushong, E.A., Ventura, P.B., Chung, W.S., Zhou, L., Cahoy, J.D., Daneman, R., Zong, H., Ellisman, M.H., et al. (2011). Development of a method for the purification and culture of rodent astrocytes. Neuron 71, 799–811.

Franchi, L., Warner, N., Viani, K., and Nunez, G. (2009). Function of Nod-like receptors in microbial recognition and host defense. Immunol Rev 227, 106–128.

Fridh, V., and Rittinger, K. (2012). The tandem CARDs of NOD2: intramolecular interactions and recognition of RIP2. PLoS One 7, e34375.

Garcia, P., Jurgens-Wemheuer, W., Uriarte Huarte, O., Michelucci, A., Masuch, A., Brioschi, S., Weihofen, A., Koncina, E., Coowar, D., Heurtaux, T., et al. (2022). Neurodegeneration and neuroinflammation are linked, but independent of alpha-synuclein inclusions, in a seeding/spreading mouse model of Parkinson’s disease. Glia 70, 935–960.

George, S., Rey, N.L., Tyson, T., Esquibel, C., Meyerdirk, L., Schulz, E., Pierce, S., Burmeister, A.R., Madaj, Z., Steiner, J.A., et al. (2019). Microglia affect alpha-synuclein cell-to-cell transfer in a mouse model of Parkinson’s disease. Mol Neurodegener 14, 34.

Gong, Q., Long, Z., Zhong, F.L., Teo, D.E.T., Jin, Y., Yin, Z., Boo, Z.Z., Zhang, Y., Zhang, J., Yang, R., et al. (2018). Structural basis of RIP2 activation and signaling. Nat Commun 9, 4993.

Gu, H., Yang, X., Mao, X., Xu, E., Qi, C., Wang, H., Brahmachari, S., York, B., Sriparna, M., Li, A., et al. (2021). Lymphocyte Activation Gene 3 (Lag3) Contributes to alpha-Synucleinopathy in alpha-Synuclein Transgenic Mice. Front Cell Neurosci 15, 656426.

Guo, J.L., and Lee, V.M. (2014). Cell-to-cell transmission of pathogenic proteins in neurodegenerative diseases. Nat Med 20, 130–138.

Guttenplan, K.A., Stafford, B.K., El-Danaf, R.N., Adler, D.I., Munch, A.E., Weigel, M.K., Huberman, A.D., and Liddelow, S.A. (2020). Neurotoxic Reactive Astrocytes Drive Neuronal Death after Retinal Injury. Cell Rep 31, 107776.

Guttenplan, K.A., Weigel, M.K., Prakash, P., Wijewardhane, P.R., Hasel, P., Rufen-Blanchette, U., Munch, A.E., Blum, J.A., Fine, J., Neal, M.C., et al. (2021). Neurotoxic reactive astrocytes induce cell death via saturated lipids. Nature 599, 102–107.

Hanson, E.K., and Whelan, R.J. (2023). Application of the Nicoya OpenSPR to Studies of Biomolecular Binding: A Review of the Literature from 2016 to 2022. Sensors (Basel) 23.

Heneka, M.T., Kummer, M.P., and Latz, E. (2014). Innate immune activation in neurodegenerative disease. Nat Rev Immunol 14, 463–477.

Hirsch, E.C., and Hunot, S. (2009). Neuroinflammation in Parkinson’s disease: a target for neuroprotection? Lancet Neurol 8, 382–397.

Hirsch, E.C., and Standaert, D.G. (2021). Ten Unsolved Questions About Neuroinflammation in Parkinson’s Disease. Mov Disord 36, 16–24.

Hjerpe, R., Aillet, F., Lopitz-Otsoa, F., Lang, V., England, P., and Rodriguez, M.S. (2009). Efficient protection and isolation of ubiquitylated proteins using tandem ubiquitin-binding entities. EMBO Rep 10, 1250–1258.

Joshi, A.U., Minhas, P.S., Liddelow, S.A., Haileselassie, B., Andreasson, K.I., Dorn, G.W., 2nd, and Mochly-Rosen, D. (2019). Fragmented mitochondria released from microglia trigger A1 astrocytic response and propagate inflammatory neurodegeneration. Nat Neurosci *22*, 1635-1648.

Kam, T.I., Hinkle, J.T., Dawson, T.M., and Dawson, V.L. (2020). Microglia and astrocyte dysfunction in parkinson’s disease. Neurobiol Dis 144, 105028.

Kam, T.I., Mao, X., Park, H., Chou, S.C., Karuppagounder, S.S., Umanah, G.E., Yun, S.P., Brahmachari, S., Panicker, N., Chen, R., et al. (2018). Poly(ADP-ribose) drives pathologic alpha-synuclein neurodegeneration in Parkinson’s disease. Science 362.

Kang, U.J., Boehme, A.K., Fairfoul, G., Shahnawaz, M., Ma, T.C., Hutten, S.J., Green, A., and Soto, C. (2019). Comparative study of cerebrospinal fluid alpha-synuclein seeding aggregation assays for diagnosis of Parkinson’s disease. Mov Disord 34, 536–544.

Kofler, J., and Wiley, C.A. (2011). Microglia: key innate immune cells of the brain. Toxicol Pathol 39, 103–114.

Kwon, S.H., Kim, S., Park, A.Y., Lee, S., Gadhe, C.G., Seo, B.A., Park, J.S., Jo, S., Oh, Y., Kweon, S.H., et al. (2021). A Novel, Selective c-Abl Inhibitor, Compound 5, Prevents Neurodegeneration in Parkinson’s Disease. J Med Chem *64*, 15091-15110.

Lagalwar, S. (2022). Mechanisms of tunneling nanotube-based propagation of neurodegenerative disease proteins. Front Mol Neurosci 15, 957067.

Liddelow, S.A., Guttenplan, K.A., Clarke, L.E., Bennett, F.C., Bohlen, C.J., Schirmer, L., Bennett, M.L., Munch, A.E., Chung, W.S., Peterson, T.C., et al. (2017). Neurotoxic reactive astrocytes are induced by activated microglia. Nature 541, 481–487.

Lim, S., Kim, H.J., Kim, D.K., and Lee, S.J. (2018). Non-cell-autonomous actions of alpha-synuclein: Implications in glial synucleinopathies. Prog Neurobiol 169, 158–171.

Lopitz-Otsoa, F., Rodriguez-Suarez, E., Aillet, F., Casado-Vela, J., Lang, V., Matthiesen, R., Elortza, F., and Rodriguez, M.S. (2012). Integrative analysis of the ubiquitin proteome isolated using Tandem Ubiquitin Binding Entities (TUBEs). J Proteomics 75, 2998–3014.

Luk, K.C., Kehm, V., Carroll, J., Zhang, B., O’Brien, P., Trojanowski, J.Q., and Lee, V.M. (2012a). Pathological alpha-synuclein transmission initiates Parkinson-like neurodegeneration in nontransgenic mice. Science 338, 949–953.

Luk, K.C., Kehm, V.M., Zhang, B., O’Brien, P., Trojanowski, J.Q., and Lee, V.M. (2012b). Intracerebral inoculation of pathological alpha-synuclein initiates a rapidly progressive neurodegenerative alpha-synucleinopathy in mice. J Exp Med 209, 975–986.

Ma, J., Gao, J., Wang, J., and Xie, A. (2019). Prion-Like Mechanisms in Parkinson’s Disease. Front Neurosci 13, 552.

Ma, S.X., and Lim, S.B. (2021). Single-Cell RNA Sequencing in Parkinson’s Disease. Biomedicines 9.

Maekawa, S., Ohto, U., Shibata, T., Miyake, K., and Shimizu, T. (2016). Crystal structure of NOD2 and its implications in human disease. Nat Commun 7, 11813.

Mao, X., Ou, M.T., Karuppagounder, S.S., Kam, T.I., Yin, X., Xiong, Y., Ge, P., Umanah, G.E., Brahmachari, S., Shin, J.H., et al. (2016). Pathological alpha-synuclein transmission initiated by binding lymphocyte-activation gene 3. Science 353.

McConnell, E.R., McClain, M.A., Ross, J., Lefew, W.R., and Shafer, T.J. (2012). Evaluation of multi-well microelectrode arrays for neurotoxicity screening using a chemical training set. Neurotoxicology 33, 1048–1057.

Michel, P.P., Hirsch, E.C., and Hunot, S. (2016). Understanding Dopaminergic Cell Death Pathways in Parkinson Disease. Neuron 90, 675–691.

Mogensen, T.H. (2009). Pathogen recognition and inflammatory signaling in innate immune defenses. Clin Microbiol Rev 22, 240–273, Table of Contents.

Nachbur, U., Stafford, C.A., Bankovacki, A., Zhan, Y., Lindqvist, L.M., Fiil, B.K., Khakham, Y., Ko, H.J., Sandow, J.J., Falk, H., et al. (2015). A RIPK2 inhibitor delays NOD signalling events yet prevents inflammatory cytokine production. Nat Commun 6, 6442.

Negri, J., Menon, V., and Young-Pearse, T.L. (2020). Assessment of Spontaneous Neuronal Activity In Vitro Using Multi-Well Multi-Electrode Arrays: Implications for Assay Development. eNeuro 7.

Ogura, Y., Inohara, N., Benito, A., Chen, F.F., Yamaoka, S., and Nunez, G. (2001). Nod2, a Nod1/Apaf-1 family member that is restricted to monocytes and activates NF-kappaB. J Biol Chem 276, 4812–4818.

Pajares, M., A, I.R., Manda, G., Bosca, L., and Cuadrado, A. (2020). Inflammation in Parkinson’s Disease: Mechanisms and Therapeutic Implications. Cells 9.

Park, H., Kam, T.I., Peng, H., Chou, S.C., Mehrabani-Tabari, A.A., Song, J.J., Yin, X., Karuppagounder, S.S., Umanah, G.K., Rao, A.V.S., et al. (2022). PAAN/MIF nuclease inhibition prevents neurodegeneration in Parkinson’s disease. Cell 185, 1943–1959 e1921.

Pellegrini, E., Desfosses, A., Wallmann, A., Schulze, W.M., Rehbein, K., Mas, P., Signor, L., Gaudon, S., Zenkeviciute, G., Hons, M., et al. (2018). RIP2 filament formation is required for NOD2 dependent NF-kappaB signalling. Nat Commun 9, 4043.

Pellegrini, E., Signor, L., Singh, S., Boeri Erba, E., and Cusack, S. (2017). Structures of the inactive and active states of RIP2 kinase inform on the mechanism of activation. PLoS One 12, e0177161.

Perry, V.H., and Teeling, J. (2013). Microglia and macrophages of the central nervous system: the contribution of microglia priming and systemic inflammation to chronic neurodegeneration. Semin Immunopathol 35, 601–612.

Philpott, D.J., Sorbara, M.T., Robertson, S.J., Croitoru, K., and Girardin, S.E. (2014). NOD proteins: regulators of inflammation in health and disease. Nat Rev Immunol 14, 9–23.

Rodriguez, L., Marano, M.M., and Tandon, A. (2018). Import and Export of Misfolded alpha-Synuclein. Front Neurosci 12, 344.

Sabbah, A., Chang, T.H., Harnack, R., Frohlich, V., Tominaga, K., Dube, P.H., Xiang, Y., and Bose, S. (2009). Activation of innate immune antiviral responses by Nod2. Nat Immunol 10, 1073–1080.

Seo, B.A., Cho, T., Lee, D.Z., Lee, J.J., Lee, B., Kim, S.W., Shin, H.S., and Kang, M.G. (2018a). LARGE, an intellectual disability-associated protein, regulates AMPA-type glutamate receptor trafficking and memory. Proc Natl Acad Sci U S A 115, 7111–7116.

Seo, B.A., Kim, D., Hwang, H., Kim, M.S., Ma, S.X., Kwon, S.H., Kweon, S.H., Wang, H., Yoo, J.M., Choi, S., et al. (2021). TRIP12 ubiquitination of glucocerebrosidase contributes to neurodegeneration in Parkinson’s disease. Neuron 109, 3758–3774 e3711.

Seo, B.A., Lee, J.H., Kim, H.M., and Kang, M.G. (2018b). Neuronal calcium channel alpha1 subunit interacts with AMPA receptor, increasing its cell surface localisation. Biochem Biophys Res Commun 498, 402–408.

Shahnawaz, M., Mukherjee, A., Pritzkow, S., Mendez, N., Rabadia, P., Liu, X., Hu, B., Schmeichel, A., Singer, W., Wu, G., et al. (2020). Discriminating alpha-synuclein strains in Parkinson’s disease and multiple system atrophy. Nature 578, 273–277.

Shahnawaz, M., Tokuda, T., Waragai, M., Mendez, N., Ishii, R., Trenkwalder, C., Mollenhauer, B., and Soto, C. (2017). Development of a Biochemical Diagnosis of Parkinson Disease by Detection of alpha-Synuclein Misfolded Aggregates in Cerebrospinal Fluid. JAMA Neurol 74, 163–172.

Smajic, S., Prada-Medina, C.A., Landoulsi, Z., Ghelfi, J., Delcambre, S., Dietrich, C., Jarazo, J., Henck, J., Balachandran, S., Pachchek, S., et al. (2022). Single-cell sequencing of human midbrain reveals glial activation and a Parkinson-specific neuronal state. Brain 145, 964–978.

Soderberg, O., Gullberg, M., Jarvius, M., Ridderstrale, K., Leuchowius, K.J., Jarvius, J., Wester, K., Hydbring, P., Bahram, F., Larsson, L.G., et al. (2006). Direct observation of individual endogenous protein complexes in situ by proximity ligation. Nat Methods 3, 995–1000.

Sterling, J.K., Adetunji, M.O., Guttha, S., Bargoud, A.R., Uyhazi, K.E., Ross, A.G., Dunaief, J.L., and Cui, Q.N. (2020). GLP-1 Receptor Agonist NLY01 Reduces Retinal Inflammation and Neuron Death Secondary to Ocular Hypertension. Cell Rep 33, 108271.

Subramaniam, S.R., and Federoff, H.J. (2017). Targeting Microglial Activation States as a Therapeutic Avenue in Parkinson’s Disease. Front Aging Neurosci 9, 176.

Takeuchi, O., and Akira, S. (2010). Pattern recognition receptors and inflammation. Cell 140, 805–820.

Tansey, M.G., Wallings, R.L., Houser, M.C., Herrick, M.K., Keating, C.E., and Joers, V. (2022). Inflammation and immune dysfunction in Parkinson disease. Nat Rev Immunol 22, 657–673.

Volpicelli-Daley, L.A., Luk, K.C., and Lee, V.M. (2014). Addition of exogenous alpha-synuclein preformed fibrils to primary neuronal cultures to seed recruitment of endogenous alpha-synuclein to Lewy body and Lewy neurite-like aggregates. Nat Protoc 9, 2135–2146.

Wang, Q., Liu, Y., and Zhou, J. (2015). Neuroinflammation in Parkinson’s disease and its potential as therapeutic target. Transl Neurodegener 4, 19.

Wong, Y.C., and Krainc, D. (2017). alpha-synuclein toxicity in neurodegeneration: mechanism and therapeutic strategies. Nat Med 23, 1–13.

Yang, Y., Yin, C., Pandey, A., Abbott, D., Sassetti, C., and Kelliher, M.A. (2007). NOD2 pathway activation by MDP or Mycobacterium tuberculosis infection involves the stable polyubiquitination of Rip2. J Biol Chem 282, 36223–36229.

Yun, S.P., Kam, T.I., Panicker, N., Kim, S., Oh, Y., Park, J.S., Kwon, S.H., Park, Y.J., Karuppagounder, S.S., Park, H., et al. (2018). Block of A1 astrocyte conversion by microglia is neuroprotective in models of Parkinson’s disease. Nat Med 24, 931–938.

Zamanian, J.L., Xu, L., Foo, L.C., Nouri, N., Zhou, L., Giffard, R.G., and Barres, B.A. (2012). Genomic analysis of reactive astrogliosis. J Neurosci 32, 6391–6410.

Zhang, Q., Duan, Q., Gao, Y., He, P., Huang, R., Huang, H., Li, Y., Ma, G., Zhang, Y., Nie, K., et al. (2023a). Cerebral Microvascular Injury Induced by Lag3-Dependent alpha-Synuclein Fibril Endocytosis Exacerbates Cognitive Impairment in a Mouse Model of alpha-Synucleinopathies. Adv Sci (Weinh) 10, e2301903.

Zhang, S., Li, J., Xu, Q., Xia, W., Tao, Y., Shi, C., Li, D., Xiang, S., and Liu, C. (2023b). Conformational Dynamics of an alpha-Synuclein Fibril upon Receptor Binding Revealed by Insensitive Nuclei Enhanced by Polarization Transfer-Based Solid-State Nuclear Magnetic Resonance and Cryo-Electron Microscopy. J Am Chem Soc 145, 4473–4484.

Zhang, S., Liu, Y.Q., Jia, C., Lim, Y.J., Feng, G., Xu, E., Long, H., Kimura, Y., Tao, Y., Zhao, C., et al. (2021). Mechanistic basis for receptor-mediated pathological alpha-synuclein fibril cell-to-cell transmission in Parkinson’s disease. Proc Natl Acad Sci U S A 118.

