## Supplementary Figures for "Pathologic α-Synuclein-NOD2 Interaction and RIPK2 Activation Drives Microglia-Induced Neuroinflammation in Parkinson’s Disease"

### Supplementary Figures and Legends

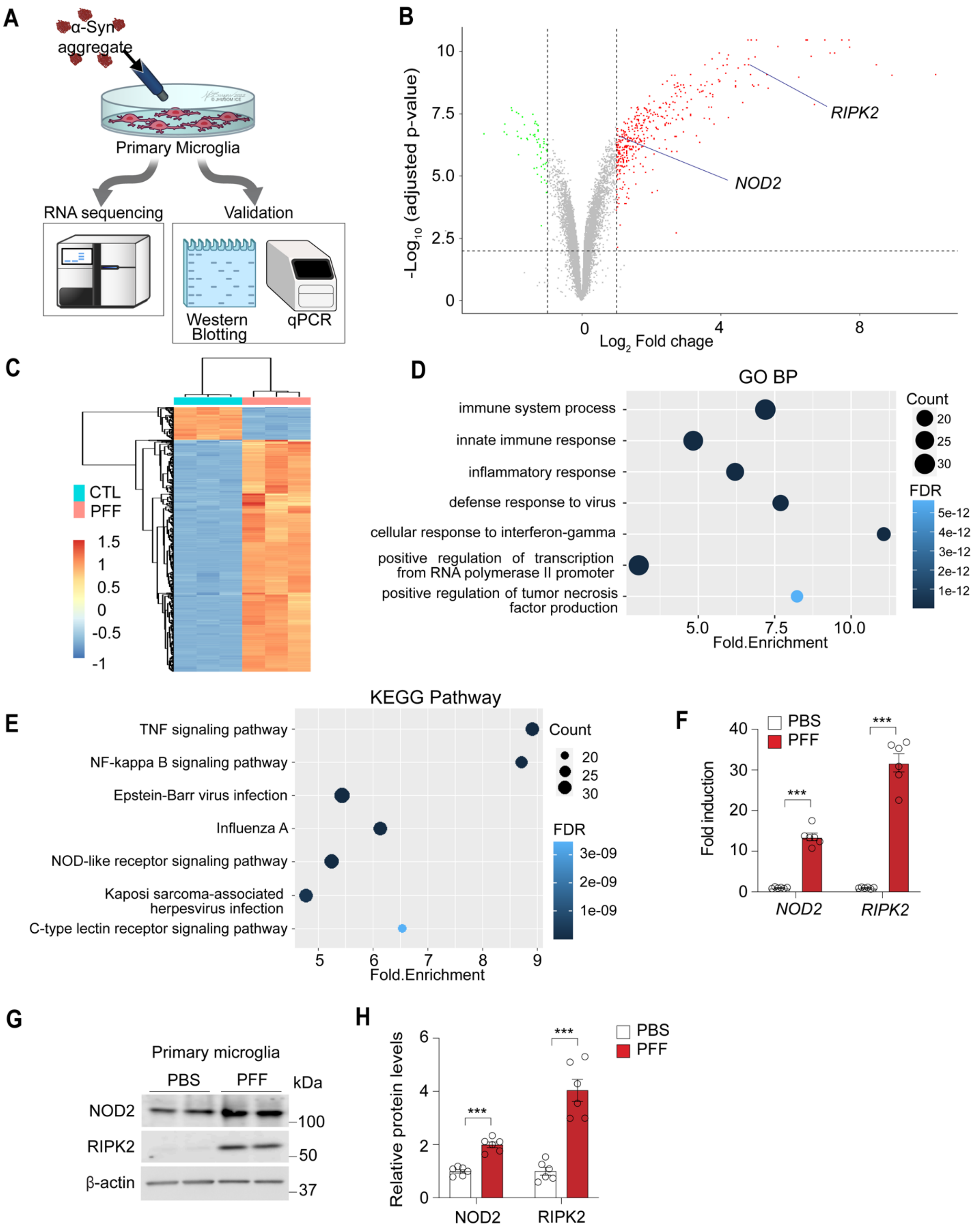

**Figure S1. *NOD2/RIPK2* transcripts and proteins are significantly upregulated in α-Syn PFF-treated microglia.**

(A) Schematic diagram depicting RNA-seq analysis of mouse primary cultured microglia treated with endotoxin-free α-Syn PFF or PBS to identify RNA quantity and sequences and qPCR and Western blotting for target validation.

(B) Volcano plots depict total transcript expressions. Red dots represent 405 up-regulated differentially expressed genes (DEGs) and green dots represent 57 down-regulated (adjusted p-value ≤ 0.01, log2(FC) ≥ 1 or log2(FC) ≤ -1). Black dots represent insignificant DEGs. The vertical dashed lines indicate a 2-fold change (log2(FC) = 1 or -1) and the horizontal dashed line indicates adjusted p-value = 0.01 (-log10 (adj. p-value) = 2). Notably, RIPK2 and NOD2 are indicated by each solid line.

(C) Hierarchical cluster heat map for displaying expression levels of 462 differentially expressed genes (DEGs) in α-Syn PFF-treated primary microglia compared to PBS-treated microglia (adjusted p-value ≤ 0.01, log2(FC) ≥ 1 or log2(FC) ≤ -1) from three comparisons. Color coding indicates the z-score values of each sample with yellow above the mean and cyan beneath (n=3, each group). Red and blue indicate up-regulation and down-regulation, respectively.

(D) Gene ontology (GO) enrichment analysis of the biological process for DEGs.

(E) Kyoto encyclopedia of genes and genomes (KEGG) enrichment analysis for DEGs. Only the top7 significant terms are shown. The size of the points represents the number of DEGs in each term and the color of point represents FDR. The x-axis of the graph represents the fold enrichment score.

(F) Quantification of increased *RIPK2* and *NOD2* mRNA levels in α-Syn PFF-treated primary microglia using qPCR (n=6, biologically independent experiments). Data are presented as mean ± SEM (***$P$ < 0.001).

(G) Immunoblots showing increased NOD2 and RIPK2 proteins in primary cultured microglia treated with α-syn PFF 5 μg/ml for 24h.

(H) Quantification of protein expression in (G) presented as a bar graph (n=6, biologically independent experiments). Data are presented as mean ± SEM (***$P$ < 0.001).

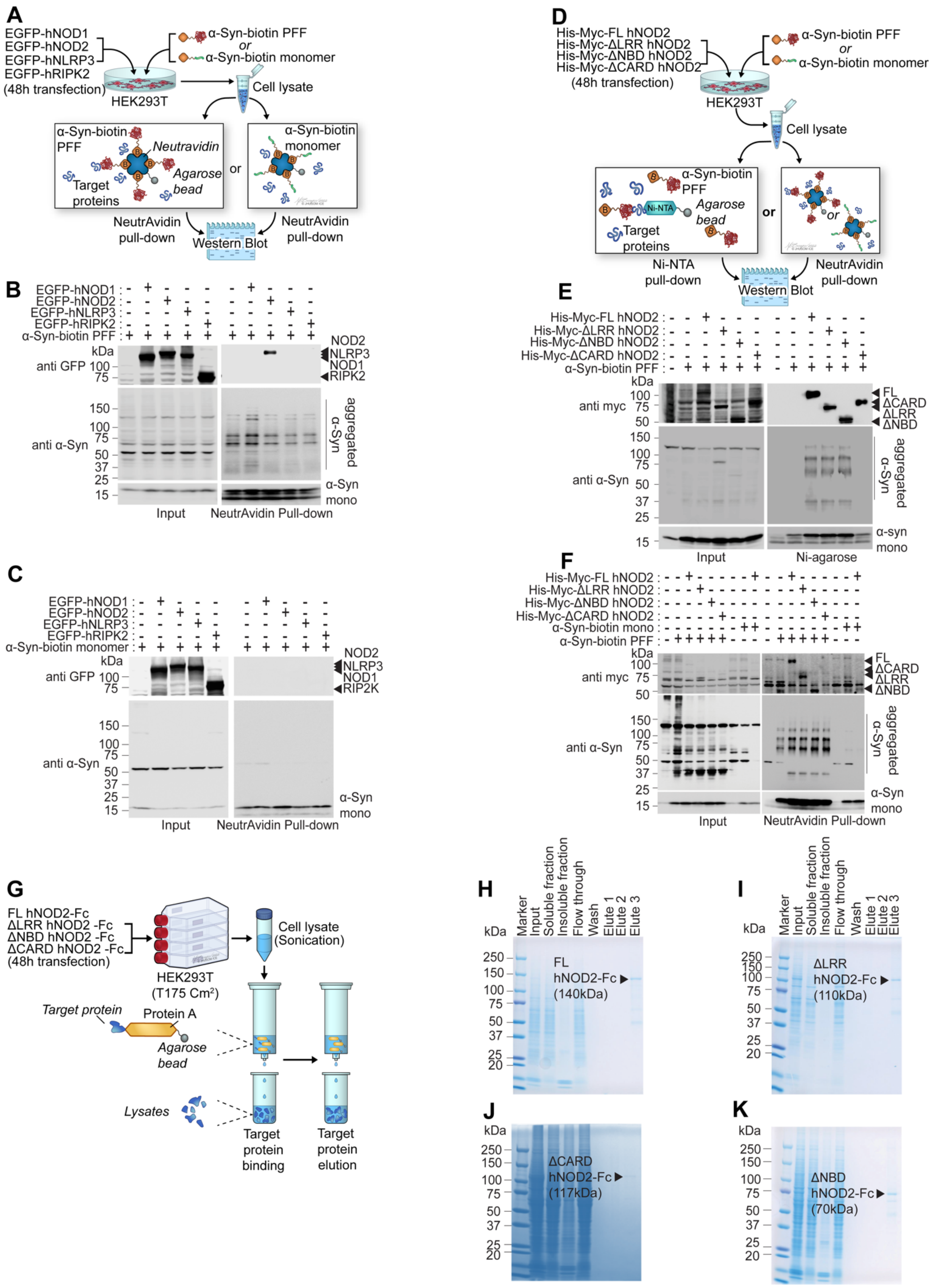
A
EGFP-hNOD1
EGFP-hNOD2
EGFP-hNLRP3
EGFP-hRIPK2
(48h transfection)
α-Syn-biotin PFF
or
α-Syn-biotin monomer
HEK293T
Cell lysate
α-Syn-biotin PFF
Neutravidin
Agarose bead
Target proteins
α-Syn-biotin monomer
NeutrAvidin pull-down
Western Blot
B
anti GFP
anti α-Syn
NOD2
NLRP3
NOD1
RIPK2
aggregated α-Syn
α-Syn mono
Input
NeutrAvidin Pull-down
C
α-Syn-biotin monomer
RIP2K
D
His-Myc-FL hNOD2
His-Myc-ΔLRR hNOD2
His-Myc-ΔNBD hNOD2
His-Myc-ΔCARD hNOD2
Ni-NTA pull-down
E
anti myc
FL
ΔCARD
ΔLRR
ΔNBD
Ni-agarose
F
α-Syn-biotin mono
G
FL hNOD2-Fc
ΔLRR hNOD2 -Fc
ΔNBD hNOD2 -Fc
ΔCARD hNOD2 -Fc
HEK293T (T175 Cm²)
Cell lysate (Sonication)
Target protein
Protein A
Lysates
Target protein binding
Target protein elution
H
Marker
Input
Soluble fraction
Insoluble fraction
Flow through
Wash
Elute 1
Elute 2
Elute 3
FL hNOD2-Fc (140kDa)
I
ΔLRR hNOD2-Fc (110kDa)
J
ΔCARD hNOD2-Fc (117kDa)
K
ΔNBD hNOD2-Fc (70kDa)

**Figure S2. Pathologic α-Syn binds to hNOD2.**

(A) Schematic of NeutrAvidin pull-down assay to confirm α-Syn PFF and hNOD2 interaction in EGFP-tagged hNOD1, -hNOD2, -hNLRP3, or -hRIPK2 transfected HEK293T cells treated with α-Syn-biotin monomer or α-Syn-biotin PFF.

(B) Representative immunoblots showing specific α-Syn PFF binding to hNOD2 (n=3, independent experiments).

(C) Representative immunoblots demonstrating no binding of α-Syn monomer to hNOD1, hNOD2, hNLRP3, or hRIPK2 (n=3, independent experiments).

(D) Schematic of in vitro reciprocal pull-down assay to verify α-Syn PFF and hNOD2 CARD domain interaction in HEK293T cells transfected with full-length and mutant hNOD2 (His-Myc-FL, His-Myc-ΔLRR, His-Myc-ΔNBD, and His-Myc-ΔCARD) and treated with α-Syn-biotin monomer or α-Syn-biotin PFF.

(E) Representative immunoblots demonstrating no binding of α-Syn PFF to ΔCARD hNOD2 using Ni-agarose pull-down (n=3, independent experiments).

(F) Representative immunoblots showing no binding of α-Syn PFF to ΔCARD hNOD2 using NeutrAvidin pull-down (n=3, independent experiments).

(G) Schematic for purification of FL hNOD2-Fc, ΔLRR hNOD2-Fc, ΔNBD hNOD2-Fc, or ΔCARD hNOD2-Fc proteins from HEK293T cells using protein A. (H-K) SDS-PAGE gels showing purified (H) FL hNOD2-Fc, (I) ΔLRR hNOD2-Fc, (J) ΔNBD hNOD2-Fc, and (K) ΔCARD hNOD2-Fc proteins for indirect ELISA.

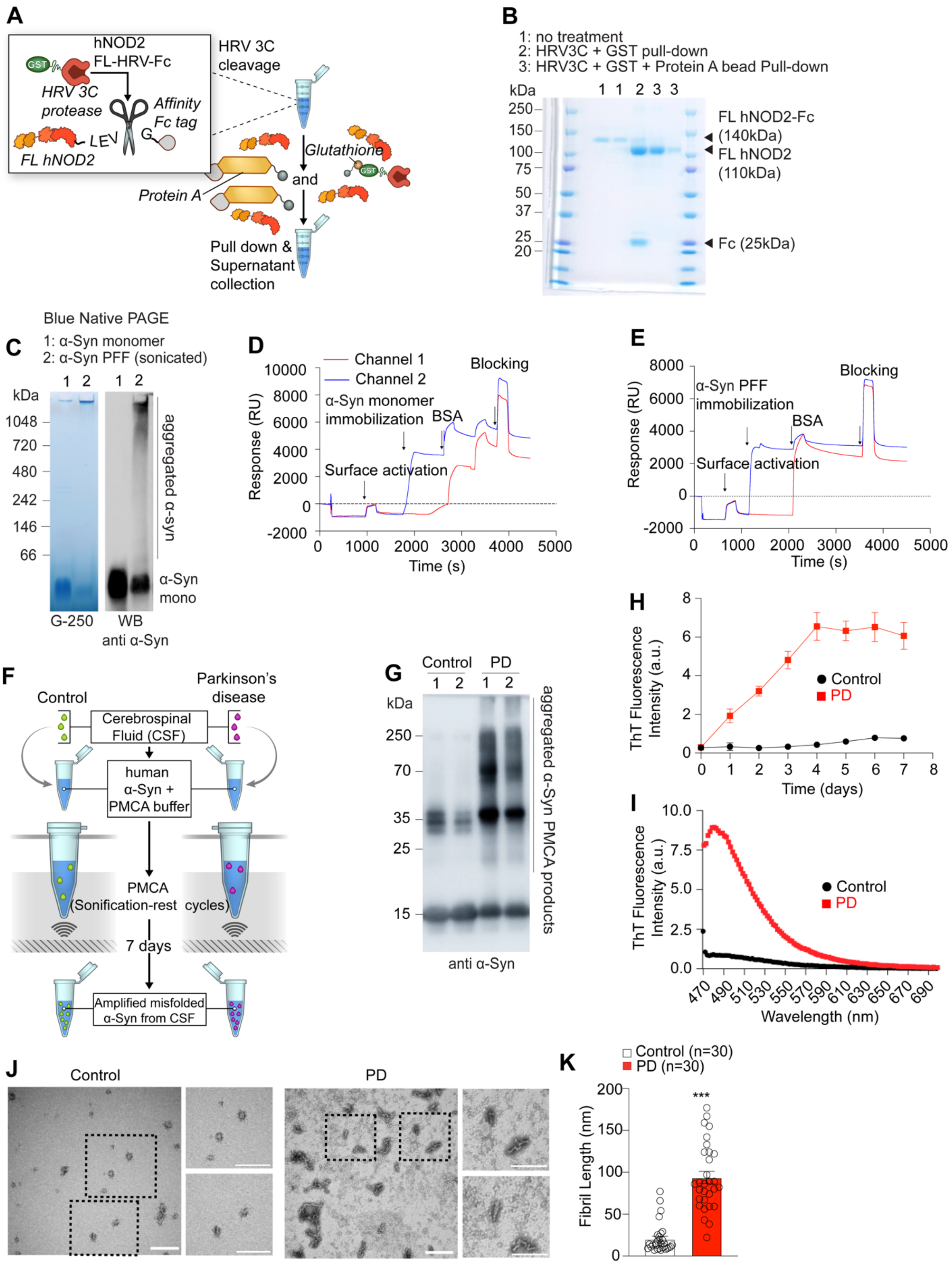
A
hNOD2
FL-HRV-Fc
GST
HRV 3C protease
Affinity Fc tag
LEV
G
FL hNOD2
HRV 3C cleavage
Protein A
Glutathione
and
Pull down & Supernatant collection
B
1: no treatment
2: HRV3C + GST pull-down
3: HRV3C + GST + Protein A bead Pull-down
kDa
1 1 2 3 3
250
150
100
75
50
37
25
20
FL hNOD2-Fc (140kDa)
FL hNOD2 (110kDa)
Fc (25kDa)
Blue Native PAGE
C
1: α-Syn monomer
2: α-Syn PFF (sonicated)
kDa
1 2 1 2
1048
720
480
242
146
66
aggregated α-syn
α-Syn mono
G-250
WB anti α-Syn
D
Channel 1
Channel 2
Blocking
α-Syn monomer immobilization
BSA
Surface activation
Response (RU)
Time (s)
10000
8000
6000
4000
2000
0
-2000
0 1000 2000 3000 4000 5000
E
8000
6000
4000
2000
0
-2000
α-Syn PFF immobilization
Blocking
BSA
Surface activation
Response (RU)
Time (s)
0 1000 2000 3000 4000 5000
F
Control
Parkinson's disease
Cerebrospinal Fluid (CSF)
human α-Syn + PMCA buffer
PMCA (Sonification-rest cycles)
7 days
Amplified misfolded α-Syn from CSF
G
Control
PD
1 2 1 2
kDa
250
70
35
25
15
aggregated α-Syn PMCA products
anti α-Syn
H
ThT Fluorescence Intensity (a.u.)
Control
PD
Time (days)
0 1 2 3 4 5 6 7 8
0 2 4 6 8
I
ThT Fluorescence Intensity (a.u.)
10.0
7.5
5.0
2.5
0.0
Control
PD
470 490 510 530 550 570 590 610 630 650 670 690
Wavelength (nm)
J
Control
PD
K
Control (n=30)
PD (n=30)
***
Fibril Length (nm)
0 50 100 150 200

**Figure S3. Preparation and characterization of FL hNOD2, α-Syn monomer, α-Syn PFF, and patient-derived α-Syn PMCA products for SPR experiments.**

(A) Schematic for hNOD2 purification with Fc tag removal using human rhinovirus (HRV) 3C protease.

(B) SDS-PAGE gel showing Fc tag-removed full-length NOD2 (FL hNOD2).

(C) Representative Blue Native (BN) PAGE gel and western blot validating α-Syn-monomer and α-Syn-PFF for SPR experiments.

(D and E) Representative sensorgrams showing the immobilization process of (D) α-Syn-monomer or (E) α-Syn-PFF onto the carboxy sensor chip for SPR experiments (n=3, biologically independent experiments).

(F) Schematic of PMCA assay for α-Syn in CSF samples from PD patients and healthy controls.

(G) Representative immunoblot of aggregated α-Syn PMCA products from PD patient group and healthy control groups.

(H) ThT binding kinetics of PMCA products from healthy controls and PD patients over time (n = 5, biologically independent experiments).

(I) ThT binding spectrum of PMCA products from healthy controls and PD patients according to wavelength (n = 5, biologically independent experiments).

(J) Representative electron micrographs (TEM) images of PMCA products from healthy controls (left) and PD patients (right). Scale bars; 200 nm.

(K) Bar graph quantifying fibril length in (U) (n=30, independent experiments). Data presented as mean ± SEM (***P < 0.001).

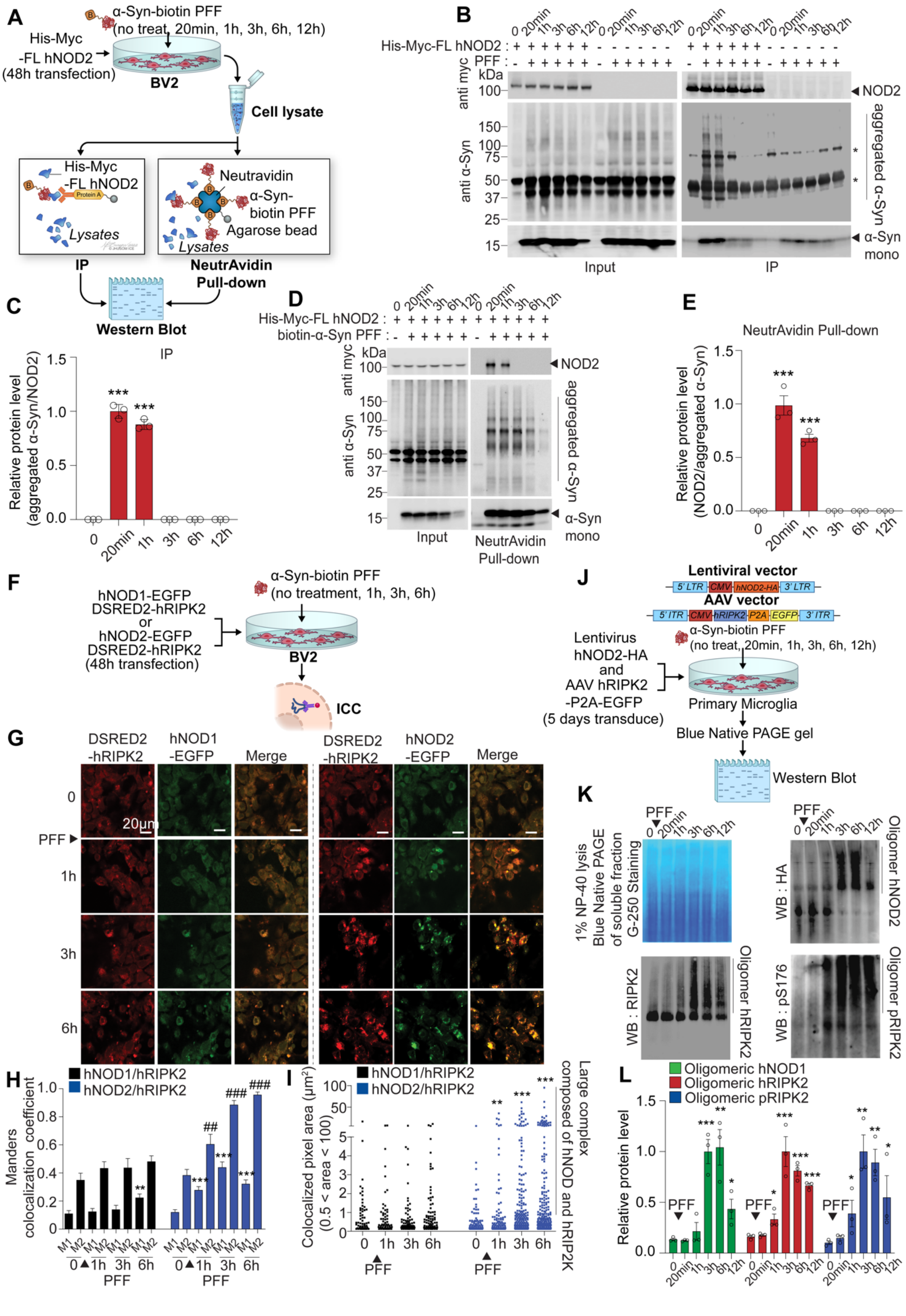
A
α-Syn-biotin PFF
(no treat, 20min, 1h, 3h, 6h, 12h)
His-Myc
-FL hNOD2
(48h transfection)
BV2
Cell lysate
His-Myc
-FL hNOD2
Protein A
Lysates
Neutravidin
α-Syn-
biotin PFF
Agarose bead
Lysates
IP
NeutrAvidin
Pull-down
Western Blot
B
His-Myc-FL hNOD2 :
PFF :
anti myc
kDa
anti α-Syn
NOD2
aggregated α-Syn
α-Syn
mono
Input
IP
C
IP
Relative protein level
(aggregated α-Syn/NOD2)
D
His-Myc-FL hNOD2 :
biotin-α-Syn PFF :
anti myc
anti α-Syn
NOD2
aggregated α-Syn
α-Syn
mono
Input
NeutrAvidin
Pull-down
E
NeutrAvidin Pull-down
Relative protein level
(NOD2/aggregated α-Syn)
F
hNOD1-EGFP
DSRED2-hRIPK2
or
hNOD2-EGFP
DSRED2-hRIPK2
(48h transfection)
α-Syn-biotin PFF
(no treatment, 1h, 3h, 6h)
BV2
ICC
G
DSRED2
-hRIPK2
hNOD1
-EGFP
Merge
DSRED2
-hRIPK2
hNOD2
-EGFP
Merge
20μm
PFF
H
hNOD1/hRIPK2
hNOD2/hRIPK2
Manders
colocalization coefficient
PFF
I
hNOD1/hRIPK2
hNOD2/hRIPK2
Colocalized pixel area (μm²)
(0.5 < area < 100)
PFF
Large complex
composed of hNOD and hRIP2K
J
Lentiviral vector
5' LTR
CMV
hNOD2-HA
3' LTR
AAV vector
5' ITR
CMV
hRIPK2
P2A
EGFP
3' ITR
α-Syn-biotin PFF
(no treat, 20min, 1h, 3h, 6h, 12h)
Lentivirus
hNOD2-HA
and
AAV hRIPK2
-P2A-EGFP
(5 days transduce)
Primary Microglia
Blue Native PAGE gel
Western Blot
K
1% NP-40 lysis
Blue Native PAGE
of soluble fraction
G-250 Staining
WB : RIPK2
Oligomer hRIPK2
WB : HA
Oligomer hNOD2
WB : pS176
Oligomer pRIPK2
L
Oligomeric hNOD1
Oligomeric hRIPK2
Oligomeric pRIPK2
Relative protein level
PFF

**Figure S4. α-Syn PFF binding to hNOD2 induces formation of microglia hNOD2 and hRIPK2 signalosome.**

(A) Schematic depicting in vitro reciprocal pull-down assay to confirm time-dependent α-Syn PFF and hNOD2 protein interaction in BV2 cells transfected with His-Myc-FL hNOD2 plasmid and treated with α-Syn-biotin PFF.

(B) Immunoblots showing that α-syn PFF interact with hNOD2 up to 1 hour and sequentially dissociate using co-immunoprecipitation assay

(C) Quantification of aggregated α-Syn of NOD2 protein expression in (B) presented as a bar graph (n=3, biologically independent experiments). Data are presented as mean ± SEM (****P* < 0.001).

(D) Immunoblots showing that α-syn PFF interact with hNOD2 up to 1 hour and sequentially dissociate using NeutrAvidin pull-down assay (n=3, biologically independent experiment).

(E) Quantification of NOD2 of aggregated α-Syn protein expression in (D) presented as a bar graph (n=3, biologically independent experiments). Data are presented as mean ± SEM (****P* < 0.001).

(F) Schematic for time-dependent immunocytochemistry in BV2 cells transfected with plasmids expressing hNOD1-EGFP and DSRED2-hRIPK2 or hNOD2-EGFP and DSRED2-hRIPK2 and treated with α-Syn PFF.

(G) Confocal images demonstrating significant hNOD2-hRIPK2 complex formation induced by α-Syn PFF in hNOD2-EGFP/DSRED2-hRIPK2 BV2 cells (right panel) and the absence of complex formation in hNOD1-EGFP/DSRED2-hRIPK2 BV2 cells (left panel).

(H) Colocalization coefficients in G, quantified as a bar graph, depict the interaction between hNOD2 and hRIPK2 proteins using Manders' coefficients: M1 (green) and M2 (red). M1 indicates the overlap with green pixels as the denominator, and M2 as the denominator for red pixels (n=13 cells, biologically independent experiments). Data are presented as mean ± SEM. For comparisons with the α-syn PFF-untreated group in M1: **P < 0.01, ***P < 0.001. For comparisons with the α-syn PFF-untreated group in M2: $^{##}$P < 0.01, $^{###}$P < 0.001.

(I) Quantification of colocalized area in G is presented as a plotted graph. The study demonstrated the progressive enlargement of complexes over time, determined by quantifying the overlapping area of fluorescence signals (green and red) above a specific threshold (n=13 cells, independent biological experiments). Data are presented as mean ± SEM (***P* < 0.01, ****P* < 0.001).

(J) Schematic diagram illustrating viral vectors expressing hNOD2-HA or hRIPK2. Immunoblotting after blue native PAGE of soluble fraction in hNOD2-HA/hRIPK2-expressing microglia stimulated with α-Syn PFF for 20 min, 1h, 3h, 6h, and 12h.

(K) Immunoblots showing hNOD2/hRIPK2 oligomerization with increasing hRIPK2 phosphorylation by α-syn PFF treatment (n=3, independent experiments).

(L) Quantification of oligomeric protein expression in (K) presented as a bar graph (n=3, biologically independent experiments). Data are presented as mean ± SEM (*$P < 0.05$, **$P < 0.01$, ***$P < 0.001$).

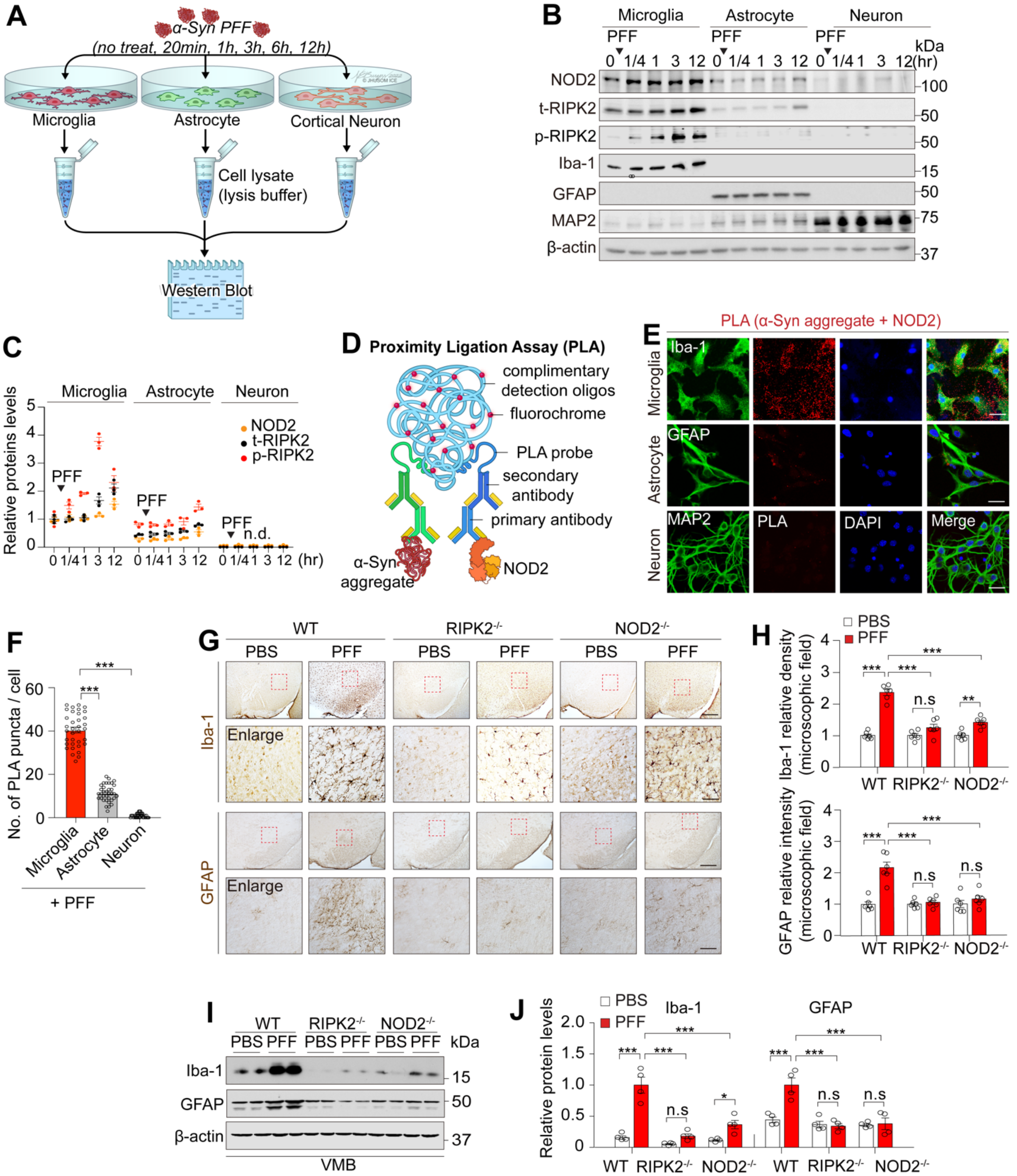


**Figure S5. Microglial NOD2 or RIPK2 deficiency reduces neuroinflammation in α-Syn PFF in mice.**

(A) Illustration showing western blotting to investigate NOD2/RIPK2 activation in mouse primary microglia, astrocytes, and cortical neurons treated with α-Syn PFF at 5 μg/ml for 20 min, 1 h, 3 h, 6 h, or 12 h.

(B) Immunoblot assessment of NOD2, t-RIPK2, and pS176 RIPK2 in a time-dependent manner in α-Syn PFF-treated primary microglia, astrocytes, or cortical neurons.

(C) Dot graph quantifying protein levels in G (n=3, biologically independent experiments).

(D) Schematic diagram depicting PLA for detecting the interaction between pathologic α-Syn aggregates and endogenous NOD2 protein in α-Syn PFF-treated mouse primary microglia, astrocytes, and cortical neurons.

(E) Confocal images showing PLA-positive signals (red dots) in α-Syn PFF-treated microglia, astrocytes, and cortical neurons.

(F) Bar graph quantifying the number of red dots per cell in E (n=35 cells, biologically independent experiments). Data are presented as mean ± SEM (***$P < 0.001$).

(G) Photomicrographs of coronal mesencephalon sections of α-Syn PFF-induced microglia and astrocytic activation in WT versus RIPK2$^{-/-}$ or NOD2$^{-/-}$ mice. Scale bars: 100 μm (upper panels) and 25 μm (lower panels).

(H) Bar graph depicting the quantification of Iba-1 relative density and GFAP relative intensity in P. (n=6, per group). Data are represented as mean ± SEM (**$P < 0.01$, ***$P < 0.001$).

(I) Representative immunoblots of α-Syn PFF-induced microglia and astrocytic activation in WT versus RIPK2$^{-/-}$ or NOD2$^{-/-}$ mice.

(J) Bar graph presenting the quantification of protein levels in R. Data are shown as mean ± SEM (n.s; not significant, *$P < 0.05$, ***$P < 0.001$).

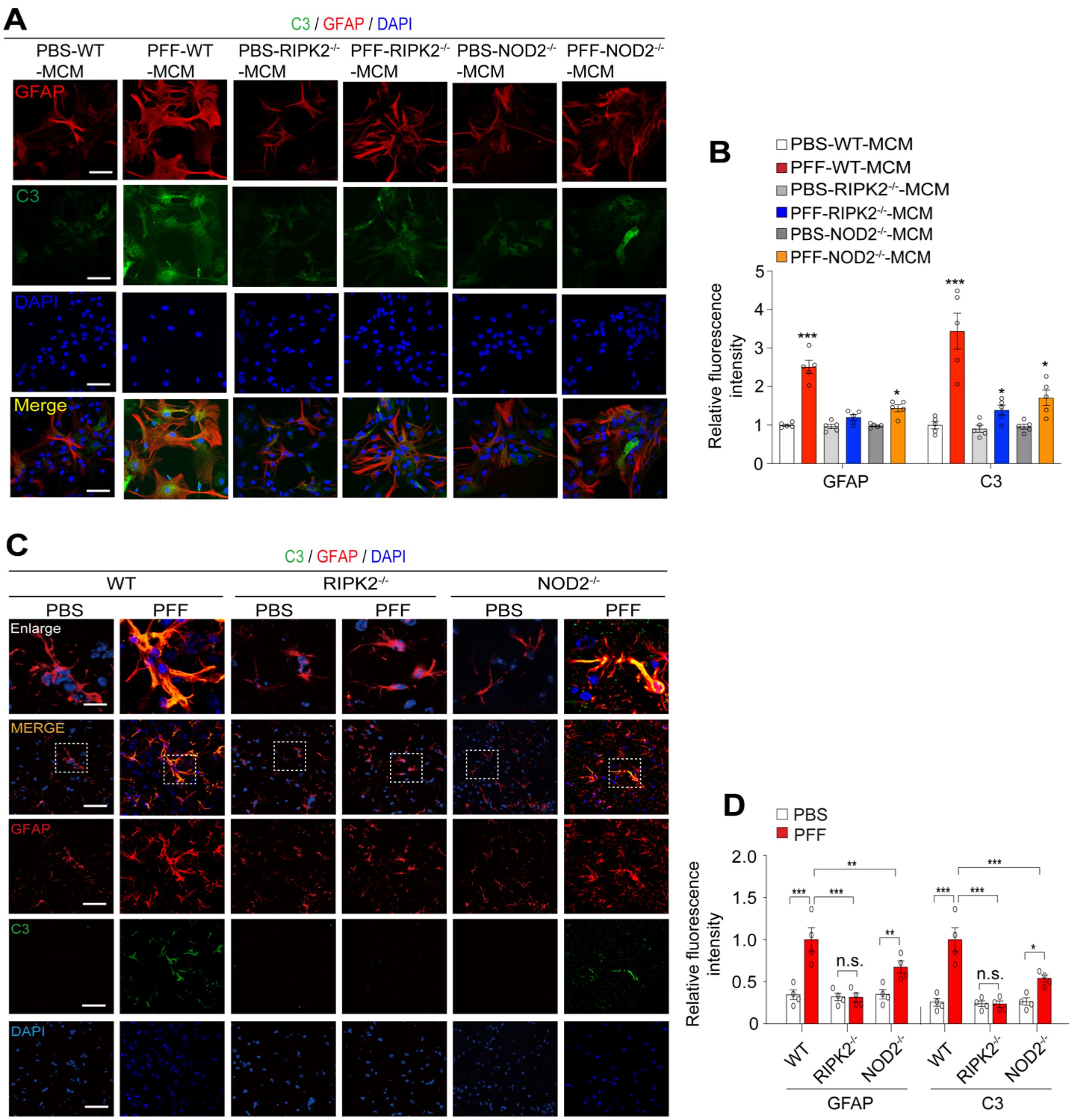


**Figure S6. NOD2 or RIPK2 deletion suppresses α-Syn PFF-induced neurotoxic reactive astrocyte formation.**

(A) Confocal images of GFAP and C3 immunoreactivity in astrocytes treated with PFF-MCM by WT, $NOD2^{-/-}$ or /$RIPK2^{-/-}$ microglia.

(B) Bar graph depicting the quantification of GFAP and C3 immunoreactivity intensity in (A) (n=5, independent experiments). Data are presented as mean ± SEM (*P < 0.05, ***P < 0.001).

(C) Confocal images of C3-positive astrocytes co-localized with GFAP immunoreactivity in α-Syn PFF-injected WT mice, $RIPK2^{-/-}$ and $NOD2^{-/-}$ mice. Specific areas (dotted line) are enlarged.

(D) Bar graph depicting the quantification of the percentage of total C3/GFAP double-positive cells in (C) (n=6, each group). Data are presented as mean ± SEM (n.s; not significant, ***$P < 0.001$).

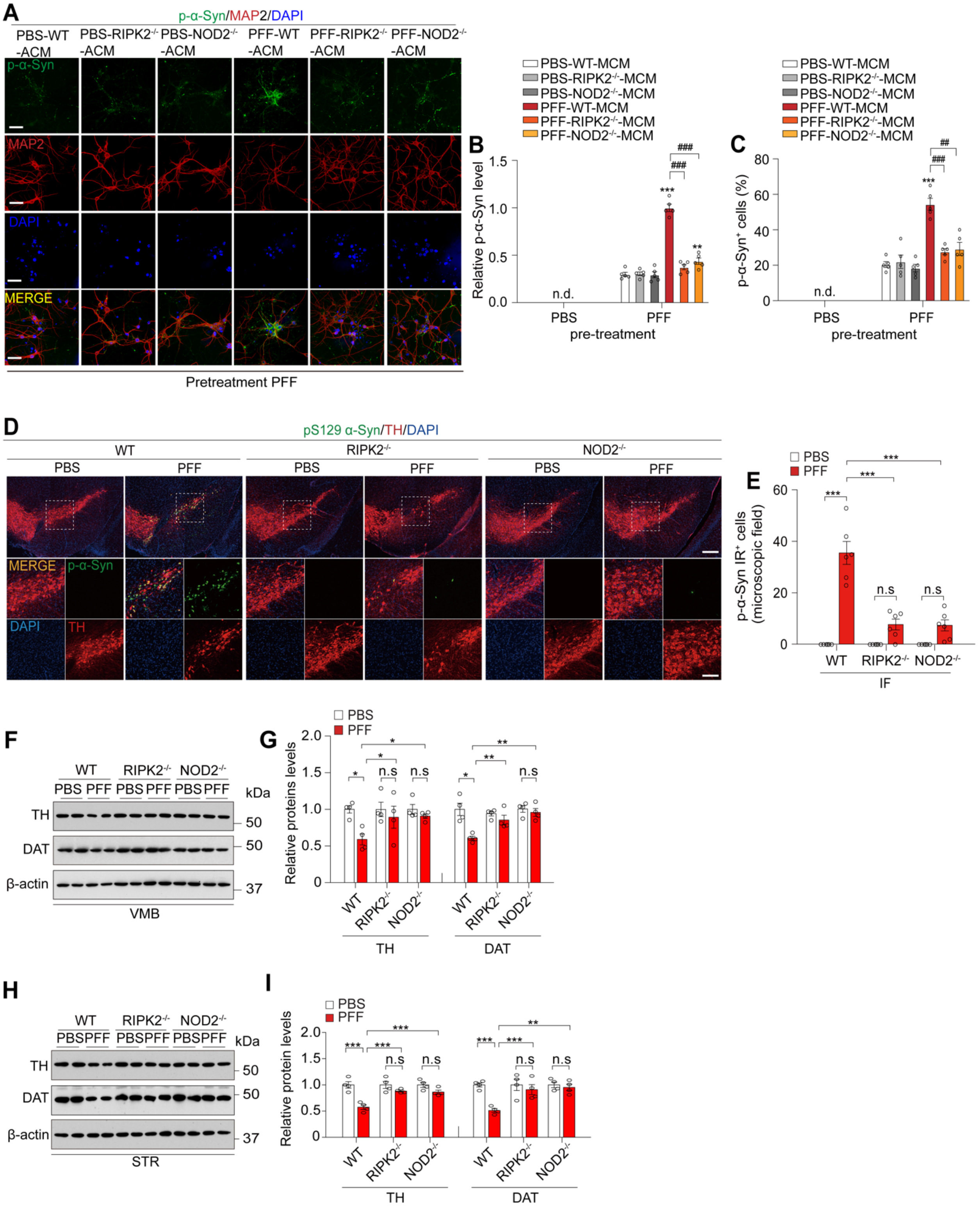


**Figure S7.** ***NOD2/RIPK2 KO*** **protects against α-Syn PFF-induced neuropathology and dopamine neuron degeneration** ***in vitro and in vivo.***

(A) Confocal images show that *RIPK2/NOD2 KO* inhibits the increase of Lewy-like pathology induced by PFF-ACM. Scale bar = 10 μm.

(B) Relative intensity of p-α-Syn immunoreactivity in MAP2-positive neurons (n=5, biologically independent experiments) in (A) shown as a bar graph. Data are presented as mean ± SEM (n.d. - no detection, for comparison to α-syn PFF-untreated group in microglia; **P < 0.01, ***P < 0.001, for comparison to α-syn PFF-treated group in WT microglia; $^{###}$P < 0.001).

(C) Percentage of p-α-Syn/MAP2 dual-positive neurons in (A) shown as a bar graph (n=5, biologically independent experiments). Data presented as mean ± SEM (n.d. - no detection, for comparison to α-syn PFF-untreated group in microglia; ***P < 0.001, for comparison to α-syn PFF-treated group in WT microglia; $^{###}$P < 0.001).

(D) Confocal images demonstrating *RIPK2/NOD2 KO* inhibition of the increase in Lewy-like pathology induced by α-Syn PFF. Double immunostaining for p-α-Syn (green) and tyrosine hydroxylase (TH, red) in the SN. White arrows indicate dopaminergic neuron loss. The white dashed box indicates a region of high magnification in the panels.

(E) Bar graph showing the number of p-α-Syn positive dopaminergic neurons in (D) (n=6, each group). Data presented as mean ± SEM (n.s - not significant, ***P < 0.001).

(F) Representative immunoblots showing that *RIPK2/NOD2 KO* rescues α-Syn PFF-induced reduction in TH and DAT levels in the ventral midbrain.

(G) Bar graph quantifying TH and DAT protein levels in (F) (n=4, each group). Data presented as mean ± SEM (n.s - not significant, *P < 0.05, **P < 0.01).

(H) Representative immunoblots showing that *RIPK2/NOD2 KO* rescues α-Syn PFF-induced reduction in TH and DAT levels in the striatum.

(I) Bar graph quantifying TH and DAT protein levels in (H) (n=4, each group). Data presented as mean ± SEM (n.s - not significant, **P < 0.01, ***P < 0.001).
